# Reconstructing the history and biological consequences of a plant invasion on the Galápagos islands

**DOI:** 10.1101/2020.09.26.313627

**Authors:** Matthew J.S. Gibson, María de Lourdes Torres, Yaniv Brandvain, Leonie C. Moyle

**Affiliations:** Department of Biology, Indiana University, Bloomington, Indiana, 47405; Universidad San Francisco de Quito (USFQ). Colegio de Ciencias Biológicas y Ambientales, Laboratorio de Biotecnología Vegetal. Campus Cumbayá. Quito, Ecuador; Galapagos Science Center, Universidad San Francisco de Quito and University of North Carolina at Chapel Hill, San Cristobal, Galapagos, Ecuador; Department of Plant Biology, University of Minnesota-Twin Cities, St. Paul, Minnesota, 55455

**Author notes:** Corresponding Author: Matthew J.S. Gibson. Author contributions: MJSG, MLT, & LCM designed the research; MJSG performed field work; MJSG performed molecular and bioinformatic analyses; YB designed the introgression HMM; MJSG implemented the HMM. MJSG wrote the manuscript, with input from YB, MLT, and LCM.

**Keywords:** Galápagos, colonization, hybridization, introgression, convergence

## Abstract

The introduction of non-native species into new habitats is one of the foremost risks to global biodiversity. Here, we evaluate a recent invasion of wild tomato (*Solanum pimpinellifolium*) onto the Galápagos islands from a population genomic perspective, using a large panel of novel collections from the archipelago as well as historical accessions from mainland Ecuador and Peru. We infer a recent invasion of *S. pimpinellifolium* on the islands, largely the result of a single event from central Ecuador which, despite its recency, has rapidly spread onto several islands in the Galápagos. By reconstructing patterns of local ancestry throughout the genomes of invasive plants, we uncover evidence for recent hybridization and introgression between *S. pimpinellifolium* and the closely related endemic species *Solanum cheesmaniae*. Two large introgressed regions overlap with known fruit color loci involved in carotenoid biosynthesis. Instead of red fruits, admixed individuals with endemic haplotypes at these loci have orange fruit colors that are typically characteristic of the endemic species. We therefore infer that introgression explains the observed trait convergence. Moreover, we infer roles for two independent loci in driving this pattern, and a likely history of selection favoring the repeated phenotypic transition from red to orange fruits. Together, our data reconstruct a complex history of invasion, expansion, and gene flow among wild tomatoes on the Galápagos islands. These findings provide critical data on the evolutionary importance of hybridization during colonization and its role in influencing conservation outcomes.

**Significance Statement:** The isolation and unique diversity of the Galápagos Islands provide numerous natural experiments that have enriched our understanding of evolutionary biology. Here we use population genomic sequencing to reconstruct the timing, path, and consequences of a biological invasion by wild tomato onto the Galápagos. We infer that invasive populations originated from a recent human-mediated migration event from central Ecuador. Our data also indicate that invasive populations are hybridizing with endemic populations, and that this has led to some invasive individuals adopting both fruit color genes and the fruit color characteristic of the endemic island species. Our results demonstrate how hybridization can shape patterns of trait evolution over very short time scales, and characterize genetic factors underlying invasive success.

## Introduction

The success of colonizing species depends on complex interactions between local environments and the availability of relevant genetic variation. Introduction events are often associated with strong genetic bottlenecks (Kolbe et al., 2004; Colautti et al., 2005; Golani et al., 2007) and reduced effective population sizes, features which may constrain the ability of colonizers to adapt to novel environments and compete with native biota (Lande, 1988; Lee, 2002). This suggests that biological invasions should rarely follow from introductions (Queller, 2000; Kolbe et al., 2004), yet successful invasions are nonetheless pervasive (Kolbe et al., 2004; Allendorf & Lundquist, 2003; Estoup et al., 2016; Comeault et al., 2020).

Several factors could be involved in this success. Despite intense bottlenecks, diversity could be maintained by other means, including multiple independent introductions (Facon et al., 2008; Kolbe et al., 2004) or via hybridization with congenerics present in the new habitat (Ellstrand & Schierenbeck, 2000; Lavergne & Molofsky, 2007; Reatini and Vision, 2020; Stepien et al., 2005). Of these mechanisms, hybridization might be particularly important for facilitating invasion into island habitats. Hybridization among native and introduced taxa is common on islands (Carlquist, 1974), potentially because of limitations on geographic extent, the abundance of generalist pollinators (Olesen et al., 2002), and/or frequent anthropogenic disturbance (Bertolo et al., 2012; Lin et al., 2013; Long et al., 2014; Cao et al., 2014). In addition, while the geographic isolation of insular habitats makes them hot spots for species endemism, only a small subset of continental taxa are successful in colonizing remote islands. The resulting incomplete trophic networks provide abundant ecological opportunities for invaders, including reproductive interactions between closely related species. Given these potentially complex contributing factors, describing the occurrence and consequences of invasion is critical for understanding both the dynamics of colonizing populations and for predicting conservation outcomes.

In this study, we investigate the contributions of demographic bottlenecks, single versus multiple introductions, and post-invasion hybridization, to patterns of genomic variation in populations of invasive and endemic tomato species on the Galápagos Islands. Two yellow/orange fruited tomato species are considered endemic to the islands: *Solanum cheesmaniae* (L. Riley) Fosberg [CHS] and *Solanum galapagense* S.C. Darwin and Peralta [GAL] (*SI Appendix, section S1*). Two red-fruited invasive species from continental Ecuador and Peru are now also documented on the archipelago: *Solanum pimpinellifolium* L. [PIM] and *Solanum lycopersicum* L. [LYC]— the domesticated tomato. Domesticated LYC was almost certainly introduced for agriculture (Rick, 1963). Wild species PIM was likely also introduced by early human colonizers (Darwin et al., 2003; Darwin, 2009), however the timing and source of this introduction is not known. Recent field surveys indicate substantially increased abundance of the invasive species while abundance of the two endemic species has markedly declined over the past two decades (Darwin, 2009; Nuez et al., 2004; Gibson et al., 2020), suggesting that recent demographic shifts may pose an extinction threat to the endemic species. Several factors also indicate a high potential for hybridization between native and invasive species, including overlapping habitats (Darwin, 2009; Nuez et al., 2004), similar flower morphologies (Darwin et al., 2003; Darwin, 2009; Rick et al., 1977; Vosters et al., 2014), and shared pollinators (Darwin, 2009). Moreover, all four species are closely related—having diverged less than 500 kya (Pease et al., 2016)—and all can be crossed to produce hybrids in the greenhouse (Rick, 1956; Rick & Bowman, 1961; Rick & Fobes, 1975).

Using genomic sequencing data from 174 plants (representing all four species) from the largest islands of San Cristobal, Santa Cruz, and Isabela, and a panel of 132 mainland PIM accessions from across the entire native range on continental South America, we (*i*) infer the timing, source, and number of invasions by PIM onto the Galápagos and (*ii*) evaluate evidence for post-colonization gene flow between the four tomato taxa, and its evolutionary consequences. We find that the majority of PIM originated in central Ecuador, are the product of a recent invasion, and are actively hybridizing with an endemic relative. By characterizing fine-scale local ancestry, we find that the emergence of novel orange-fruited plants—which resemble the endemic species in color—in two invasive populations can be explained by endemic introgression at distinct carotenoid loci with known phenotypic effects specifically on fruit color. Our findings reconstruct a recent path of invasion via Ecuador, provide evidence for ongoing interspecific gene flow, and suggest a history of natural selection favoring orange fruits in the island habitat.

## Results

### Sequencing and collections

Sequence data were drawn from 306 individual samples. We performed double-digest RAD (ddRAD) sequencing (using *PstI* and *EcoRI* enzymes) of 174 wild collected individuals from 13 populations of endemic and invasive tomatoes from three islands in the Galápagos archipelago: San Cristobal, Santa Cruz, and Isabela (**Figure 1**; **Table 1;** *SI Appendix, Fig. S5 and Table S1*). We complemented these data with ddRAD reads from 132 mainland PIM (*SI Appendix, Fig. S1* and *Table S2*), previously sequenced in Gibson & Moyle (2020) using the same enzymes. We recovered 18573 high quality RAD loci, each sequenced to an average of 61.4X (s.d. = 35X) in 80% of all 306 samples (*SI Appendix, Table S3 and Table S4*). Average insert size was 192 bp (s.d. = 51.7) after adapter and quality trimming. After filtering for depth (> 8 reads), 11297 SNPs were retained. After filtering for LD (r^2^ < 0.7), 5767 SNPs were retained. Refer to *SI Appendix, Table S5* for a summary of each filtering step and the analyses for which each dataset was used.

**Table 1:** Diversity statistics for focal population samples (H = observed heterozygosity; π = genome wide nucleotide diversity).

| Taxa | Population | Island | Endemic | Segregating Sites | H | $\pi$ |
| --- | --- | --- | --- | --- | --- | --- |
| PIM | Peru | N | N | 32820 | 0.00025 | 0.00094 |
|  | Ecuador | N | N | 21773 | 0.00033 | 0.00130 |
|  | MG105 | Y | N | 1520 | 0.00011 | 0.00009 |
|  | MG107 | Y | N | 4434 | 0.0002 | 0.00034 |
|  | MG111 | Y | N | 2562 | 0.00012 | 0.00010 |
|  | MG113 | Y | N | 3763 | 0.00015 | 0.00029 |
|  | MG114 | Y | N | 4730 | 0.00016 | 0.00023 |
|  | MG116 | Y | N | 4929 | 0.00024 | 0.00024 |
|  | MG117 | Y | N | 4776 | 0.00016 | 0.00028 |
| CHS | MG115 | Y | Y | 8540 | 0.00023 | 0.00045 |
|  | MG120C | Y | Y | 854 | 0.0001 | 0.00008 |
| GAL | MG120G | Y | Y | 3282 | 0.00013 | 0.00014 |
| GALxCHS | MG120GC | Y | Y | 2857 | 0.00023 | 0.00026 |
| LYC | MG125 | Y | N | 7219 | 0.00035 | 0.00063 |
|  | MG126 | Y | N | 4567 | 0.00018 | 0.00052 |

**Figure 1:**
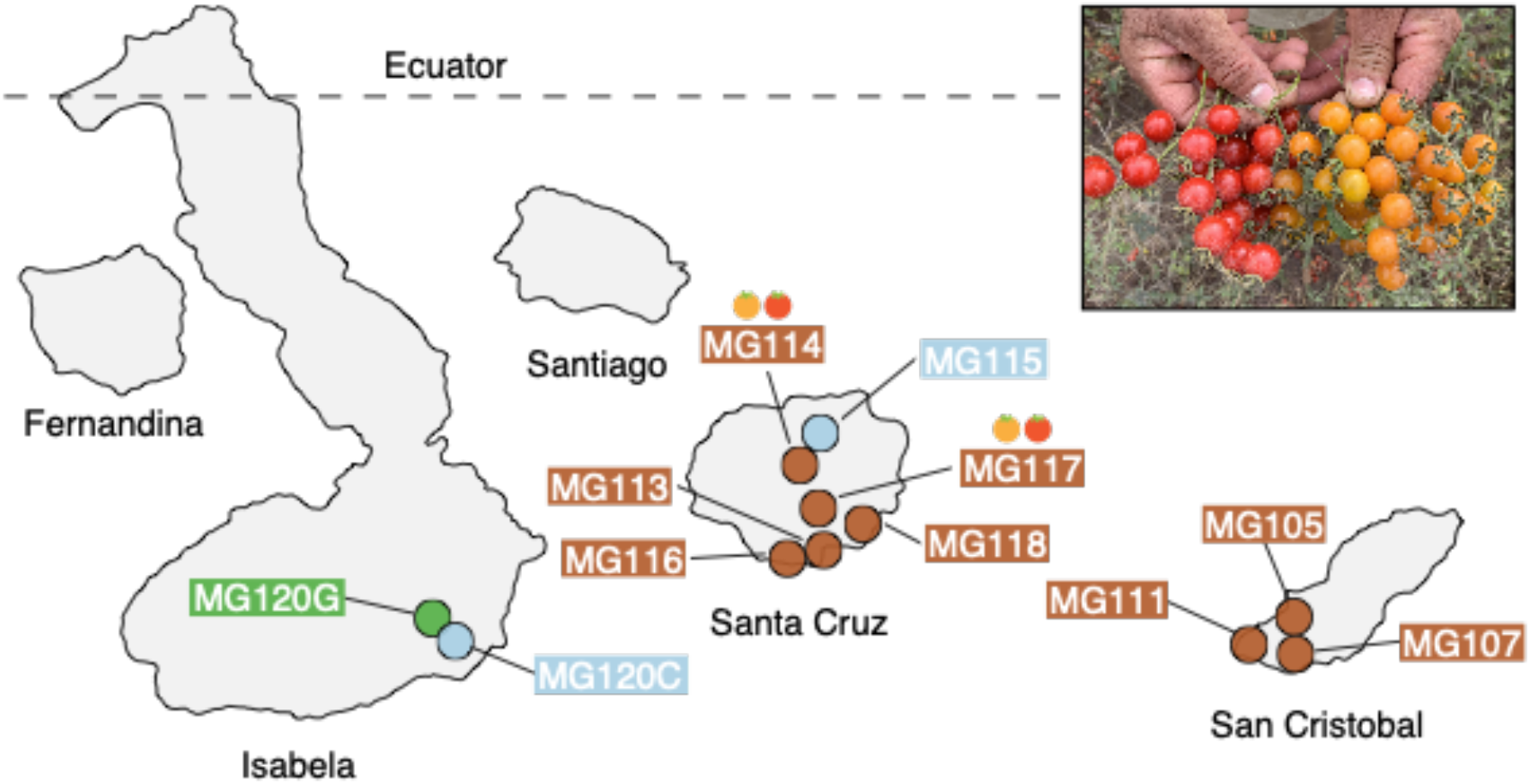
Geographic distribution of focal sampling sites on the Galápagos Islands. *Inset*: Photograph of polymorphic (red/orange) PIM fruits representative of populations MG114 and MG117. For simplicity, LYC populations as well as sampling sites with < 8 individuals are not included here. Refer to Table S1 for a full list of collection localities and sample sizes.

### Genetic data support an Ecuadorian origin for most invasive populations

Using our ddRAD sequencing data for Galápagos and continental PIM, we analyzed population genetic signatures of colonization and characterized the origin and path of invasion into the archipelago. Nucleotide diversity (π in 100kb overlapping windows; **Figure 2C**) was reduced on average 6.6-fold in island populations relative to mainland accessions (**Table 1**), a pattern consistent with population genetic expectations following colonization.

**Figure 2:**
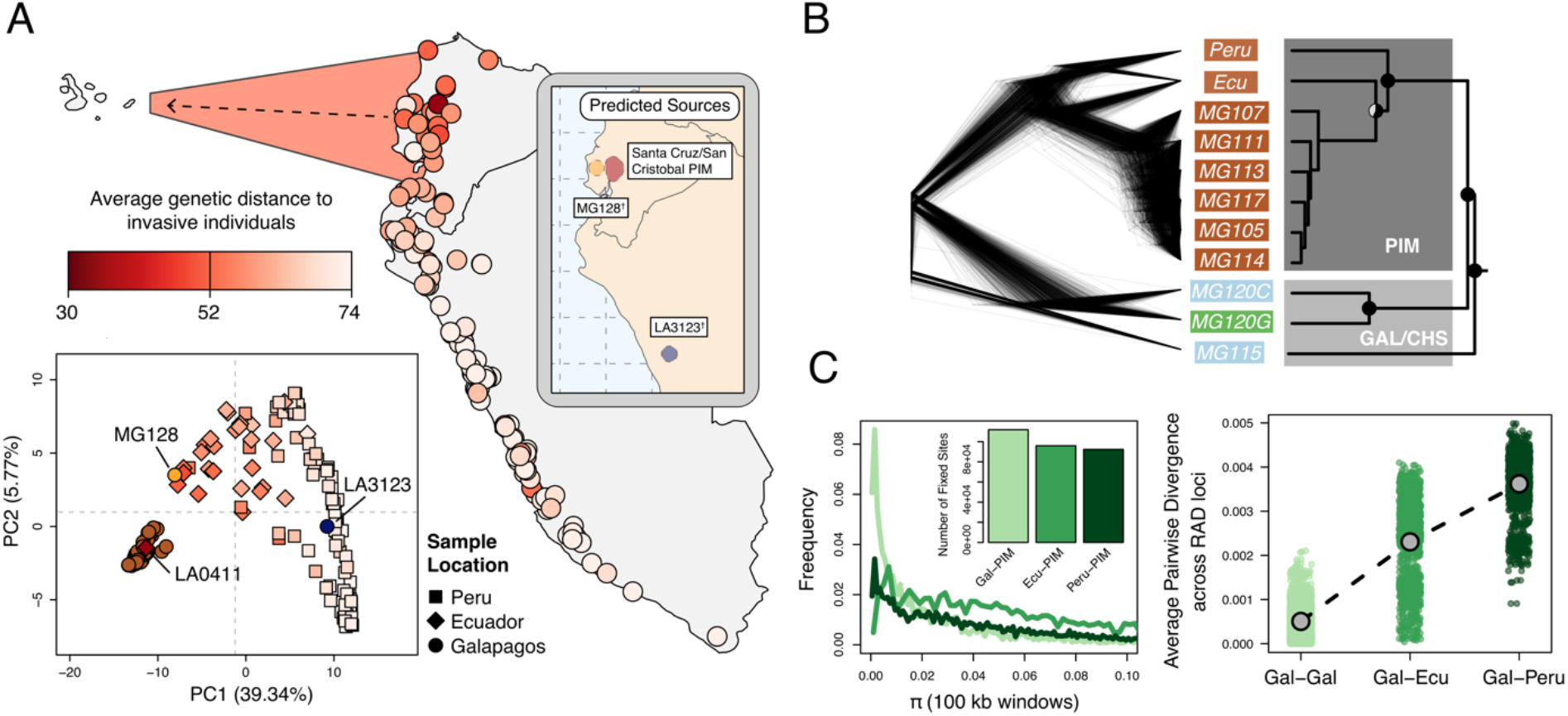
Galápagos PIM is the result of a recent invasion from Ecuador. (A) *Map:* average genetic distance between Galápagos-PIM collections and each of the 140 mainland accessions. *Plot*: multi-locus PCA. Squares, diamonds, and circles indicate Peruvian, Ecuadorian, and Galápagos collections, respectively. *Inset:* Predicted continental origins for Galápagos PIM collections. Colors are same as shown in the multi-locus PCA (^†^Exact locations vary substantially between runs. Results from a single run are shown). (B) Maximum likelihood relationships among focal populations calculated with *Treemix* (allowing no migration). *Left*: inferred trees of 1000 resampled datasets (500 SNPs, with replacement). *Right*: consensus topology. All trees were rate smoothed (λ = 1). (C) Diversity and divergence metrics. *Left*: nucleotide diversity (π) calculated for Galápagos-PIM, Ecuador-PIM, and Peru PIM in overlapping 100kb windows. Invariant windows (π = 0) are truncated and are instead shown in the inset bar plot. *Right*: average pairwise sequence divergence for three PIM comparisons: Gal x Gal, Gal x Ecu, and Gal x Peru. Each point represents a comparison between individuals, averaged over all loci.

Genetic variation in the native (mainland) range of PIM is highly geographically structured (Gibson & Moyle, 2020; *SI Appendix, Fig. S1*), allowing us to infer a putative origin of PIM lineages invasive on the Galápagos. To do so, we estimated genome-wide patterns of relatedness between invasive and mainland individuals using several methods. A rate-smoothed maximum likelihood tree constructed in *Treemix* (Pickrell & Pritchard, 2012) identified Galápagos PIM as monophyletic, and clearly separated island and non-island clades (**Figure 2B**). In general, pairwise sequence divergence was lower in Galápagos-Ecuador comparisons (average d_xy_ = 2.3 ⨯ 10^3^) than between Galápagos-Peru comparisons (average d_xy_ = 3.6 ⨯ 10^3^; **Figure 2C**), and samples showing low genome-wide divergence were clustered in central Ecuador (similar patterns were observed using F_ST_; *SI Appendix, Table S10*).

To investigate potential source localities for invasive populations at a finer scale, we implemented the software *Locator* (Battey et al., 2020) which uses a machine learning algorithm to predict sample origins from genotype data. *Locator* predictions indicated 2 to 3 source regions for Galápagos PIM, although the exact locations varied across runs and depended on which island PIM collections were considered (*SI Appendix, Fig. S2*). Santa Cruz and San Cristobal PIM collections were predicted to have originated in central Ecuador; this result was generally consistent across runs, with the consensus being an origin near Los Rios and Guayas provinces in southcentral Ecuador (*SI Appendix, Fig. S2*). Interestingly, we also infer that one mainland accession represents a back migration from the Galápagos to Los Rios (LA0411; *SI Appendix, section S2*), further highlighting the high degree of connectivity between this region and the islands. In contrast, the remaining two samples, LA3123 (a historical collection from Santa Cruz sampled in 1991) and MG128-1 (newly sampled on Isabela), were predicted to have originated in alternative locations, with most runs supporting a Peruvian origin for LA3123 and an Ecuadorian origin for MG128-1 (*SI Appendix, Fig. S2*). The exact origin locations for these samples varied substantially across runs. In general, *Locator* predictions were consistent with the pattern of low pairwise sequence divergence between Galápagos PIM and central Ecuadorian samples, pointing to Ecuador, and perhaps central Ecuador in particular, as the source of the majority of invasive PIM populations on the Galápagos.

Together our data support 2-3 independent introductions of PIM onto the archipelago, each with variable consequences for current invasive populations: (i) a minor event from Peru [LA3123], (ii) a minor event from Ecuador [MG128-1], and (iii) a major event from central Ecuador that is responsible for nearly all sampled populations.

### Demographic reconstruction supports a recent colonization by PIM on the Galápagos

We used the allele frequency spectrum to model the demographic history of invasive populations. In particular, we evaluated two demographic models using *δaδi* (Gutenkunst et al., 2010): (i) a neutral model of constant population size and (ii) an instantaneous population bottleneck model. Since this species is self-fertile (that is, it lacks genetic self-incompatibility that is present in some wild tomato species; Rick et al., 1977) we simultaneously inferred the inbreeding coefficient (F; Blischak et al., 2020). The bottleneck model thus included 5 parameters: bottleneck population size (N_B_), final population size (N_F_), timing of the bottleneck (T_B_), timing of recovery (T_F_), and the inbreeding coefficient. To limit potential confounding effects due to population structure within PIM, we estimated the folded site frequency spectrum (SFS) of a single population (MG114, which was the most deeply sampled of our PIM populations). We also masked regions of inferred introgression (as detected by our HMM, see below) as these can spuriously inflate rare variants and thus bias the inference of a bottleneck and subsequent parameter estimation (*SI Appendix, section S6*). In our masked dataset, we observed an excess of rare variants (genome wide Tajima’s D = -0.48) more consistent with a bottleneck model (RSS_Bottle_ = 1.39; ln(L) = -18.93) than a neutral model (RSS_Neutral_ = 15.12; ln(L) = -25.89).

We used the best-fit bottleneck model to estimate the timing of the introduction, performing the *δaδi* optimization procedure described in Portik et al. (2017). We inferred a recent bottleneck occurring 284.1 generations in the past (**Table 2**). During the bottleneck, N_E_ (effective population size) was reduced by 27% relative to the ancestral reference (N_Ref_). Variance in all parameter distributions as estimated by a nonparametric bootstrap was high (*SI Appendix, Fig. S6*), and median estimates deviated slightly from the optimized estimates (**Table 2**). In all cases, the optimized parameter estimate fell within the bootstrapped 95% CIs.

**Table 2:** Demographic model estimates for PIM population MG114 inferred using *δaδi*. 95% CI values were obtained from 2,000 bootstrap replicates of the SFS. Each estimate is shown in rescaled units (rescaled by N_Ref_ for N_B_ and N_F_; and by 2N_Ref_ for T_B_ and T_F_).

| Parameter | Optimum | Bootstrap Median | 95% CI |
| --- | --- | --- | --- |
| $N_B$ | 184.69 | 286.16 | 4.49 - 1483.26 |
| $N_F$ | 2730.97 | 977.28 | 319.12 – 5333.40 |
| $T_B$ | 43.12 | 846.90 | 91.88 – 7871.41 |
| $T_F$ | 240.98 | 839.13 | 22.28 – 13591.11 |
| F | 0.24 | 0.02 | $2.18 \times 10^{-10} - 0.54$ |

**Table 2:** Demographic model estimates for PIM population MG114 inferred using $\delta a \delta i$ .
| Parameter | Optimum | Bootstrap Median | 95% CI |
| --- | --- | --- | --- |
| $N_B$ | 184.69 | 286.16 | 4.49 - 1483.26 |
| $N_F$ | 2730.97 | 977.28 | 319.12 – 5333.40 |
| $T_B$ | 43.12 | 846.90 | 91.88 – 7871.41 |
| $T_F$ | 240.98 | 839.13 | 22.28 – 13591.11 |
| $F$ | 0.24 | 0.02 | $2.18 \times 10^{-10}$ – 0.54 |

For comparison, we also model the history of an endemic CHS population (MG115) using the same framework as above. A bottleneck model again fit the data better (ln(L) = -49.91) than a neutral model (ln(L) = -73.51). We estimate the timing of the bottleneck at 3,747.39 generations in the past, approximately 13X higher than the estimate for PIM. Similarly, N_E_ during the bottleneck was reduced 34% relative to the ancestral population and subsequently increased following the bottleneck (increase of 1.7-fold relative to N_Ref_).

### Admixture analyses support the occurrence of inter- and intraspecific gene flow

The close evolutionary relationship of PIM, CHS, and GAL, their similar floral morphologies, and the presence of only a single major pollinator on the islands (*Xylocarpii darwini*; McMullen, 1999), indicate the potential for interspecific gene flow between tomato species may be high. Key morphological observations also suggest that these species may be exchanging genes (Darwin, 2009). In particular, we have previously described a novel fruit color polymorphism in two Santa Cruz PIM populations (MG114 & MG117; Gibson et al., 2020), where approximately 40% of individuals have orange instead of their ancestrally red fruits. Orange fruits are very rare in mainland PIM (TGRC passport data; www.tgrc.ucdavis.edu) but are diagnostic of the two endemic Galápagos species. Accordingly, we used multiple population genomic methods to investigate evidence of hybridization and introgression in the genomes of island plants, paying special attention to patterns of admixture in the polymorphic PIM populations.

We first examined evidence for recent (early generation) hybrids by evaluating genome-wide signatures in *fastStructure* (Raj et al., 2014) and *NewHybrids* (Anderson & Thompson, 2002). Interestingly, we find no evidence of early generation CHSxPIM hybrids in either of the polymorphic PIM populations MG114 and MG117 (**Figure 3A**). However, these analyses did detect variable levels of CHS x PIM admixture at the nearby site MG115 (**Figure 3A**), a pattern which is also reflected in principle component-space (*SI Appendix, Fig. S8*). Using *NewHybrids*, we classified 4/6 of these admixed plants as first- or second-generation hybrids (*SI Appendix, Table S6 and S8*).

**Figure 3:**
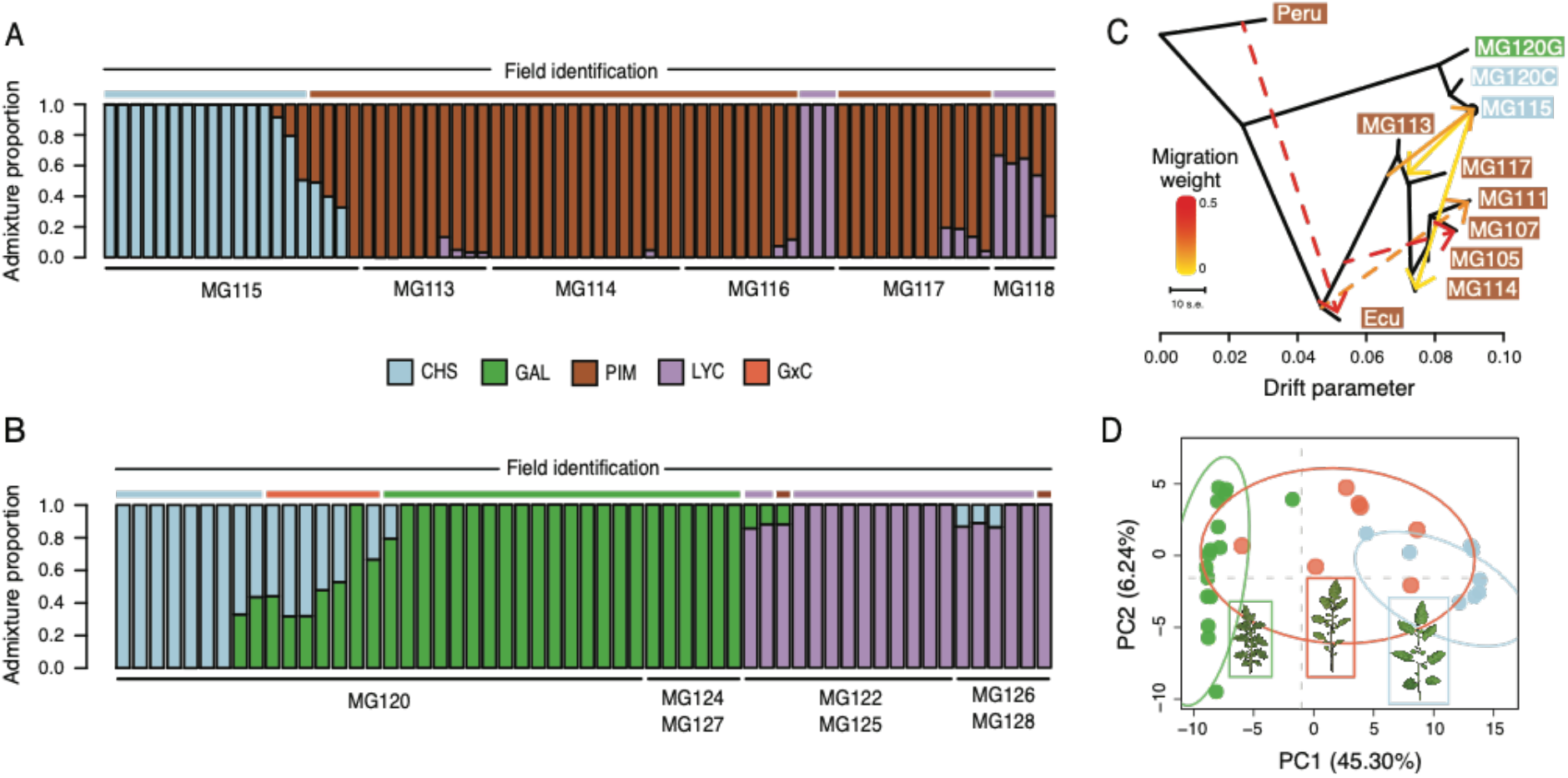
Patterns of population genetic structure and admixture on Santa Cruz and Isabela. (A) *fastStructure* inference for all Santa Cruz samples (N = 74). K = 3. (B) *fastStructure* inference for all Isabela samples (N = 57). K = 3. (C) *Treemix* analysis summary (m = 6; ln[L] = 395.08). (D) Principle components analysis for samples at site MG120, a hybrid zone between CHS and GAL.

To complement the above analyses, we employed *Treemix* (Pickrell & Pritchard, 2012). The most likely topology inferred by *Treemix* implies three separate admixture events between PIM and CHS: two cases of CHS → PIM admixture and one case of PIM → CHS admixture on Santa Cruz (**Figure 3C**). This analysis therefore indicates repeated gene flow between PIM and CHS, although how many distinct events were involved is difficult to infer given the high genomic similarity of island PIM populations and recency of the invasion (*SI Appendix, Fig. S3 and S4*). As independent support for a history of gene flow, we calculated the four taxa D statistic of Durand et al. (2011) using *Solanum pennellii* (LA3778) as an outgroup and treating PIM population MG114 and CHS (MG115) as P2 and P3, respectively. We found that D was 0.818 (s.d. = 0.028; bootstrapped P < 0.02), indicating an excess of ABBA sites and strong evidence for admixture between island PIM and CHS. D was also significant when other invasive PIM populations--Santa Cruz population MG117 or San Cristobal populations MG107 and MG105—were used as P2, indicating that the detected admixture likely predates the dispersal and differentiation between Santa Cruz and San Cristobal invasive PIM. This is consistent with inferences in the *Treemix* graph, in which admixture events between PIM and CHS involve internal branches that subtend current San Cristobal and Santa Cruz PIM populations (**Figure 3C**).

Interestingly, *fastStructure* and *Treemix* produced conflicting results regarding evidence for admixture in the polymorphic PIM populations; *Treemix* appears to support this while *fastStructure* does not. To evaluate whether this was due to differences in the detection of more subtle—and potentially older—signals of introgression, we implemented a local ancestry assignment algorithm using a Hidden Markov Model (HMM) to probe for evidence of introgression at a finer scale. Doing so, we found evidence for bidirectional gene flow between CHS and PIM (**Figure 4**; *SI Appendix, Fig. S13*), with inferred introgression being more common in the CHS → PIM direction and in PIM populations that were polymorphic for fruit color. Our HMM detected clear evidence for CHS ancestry within the polymorphic PIM populations MG114 and MG117, reflecting admixture from CHS → PIM. In contrast, inferred admixture in nonpolymorphic PIM (e.g., MG116) was much more restricted. For MG114, CHS ancestry blocks were large (average = 16,235 kb), of varying size (sd = 139.43 kb), and composed on average 3.64% of the genomes of any given MG114 plant (**Figure 4C, 4D**). Shared ancestry between MG114 and MG115 was dominated by two large CHS haplotypes segregating at moderate to high frequencies on chromosome three (40%; mean size = 51.35 Mb) and chromosome six (20%; mean size = 35.3 Mb**; Figure 4B**). The genomic distribution of CHS ancestry blocks in different individuals indicates they are not independent. For example, on chromosome 3 all but one individual (5/6) carrying the CHS haplotype had identical breakpoints, consistent with them being derived from the same hybridization (and subsequent recombination) event and/or the individuals being closely related (*SI Appendix, Table S10*). In MG117, CHS ancestry made up 4.36% (sd = 2.56%) of any given MG117 genome and average block size was 9,687.5 kb (**Figure 4C, 4D;** *SI Appendix, Table S12*). As with MG114, a large CHS haplotype on chromosome six occurs at a frequency of 0.42. This block varied substantially in size in each individual, and all were discontinuous across the chromosome (i.e., there is an implied double crossover event). Further, two individuals were heterozygous for ancestry at the downstream portion of the haplotype (*SI Appendix, Fig. S10, S11, S12*). In comparison to MG114 and MG117, MG116 showed little evidence for shared ancestry. While 3/7 individuals had inferred signals of CHS ancestry, these blocks were generally small compared to MG114 and MG117 (average = 4,450 kb; **Figure 5**) and made up a substantially smaller fraction of the total genome (average admixture proportion = 0.55%; **Figure 4D**). No large CHS haplotypes were segregating in MG116, unlike those observed in MG114 and MG117 (*SI Appendix, Table S13*).

**Figure 4:**
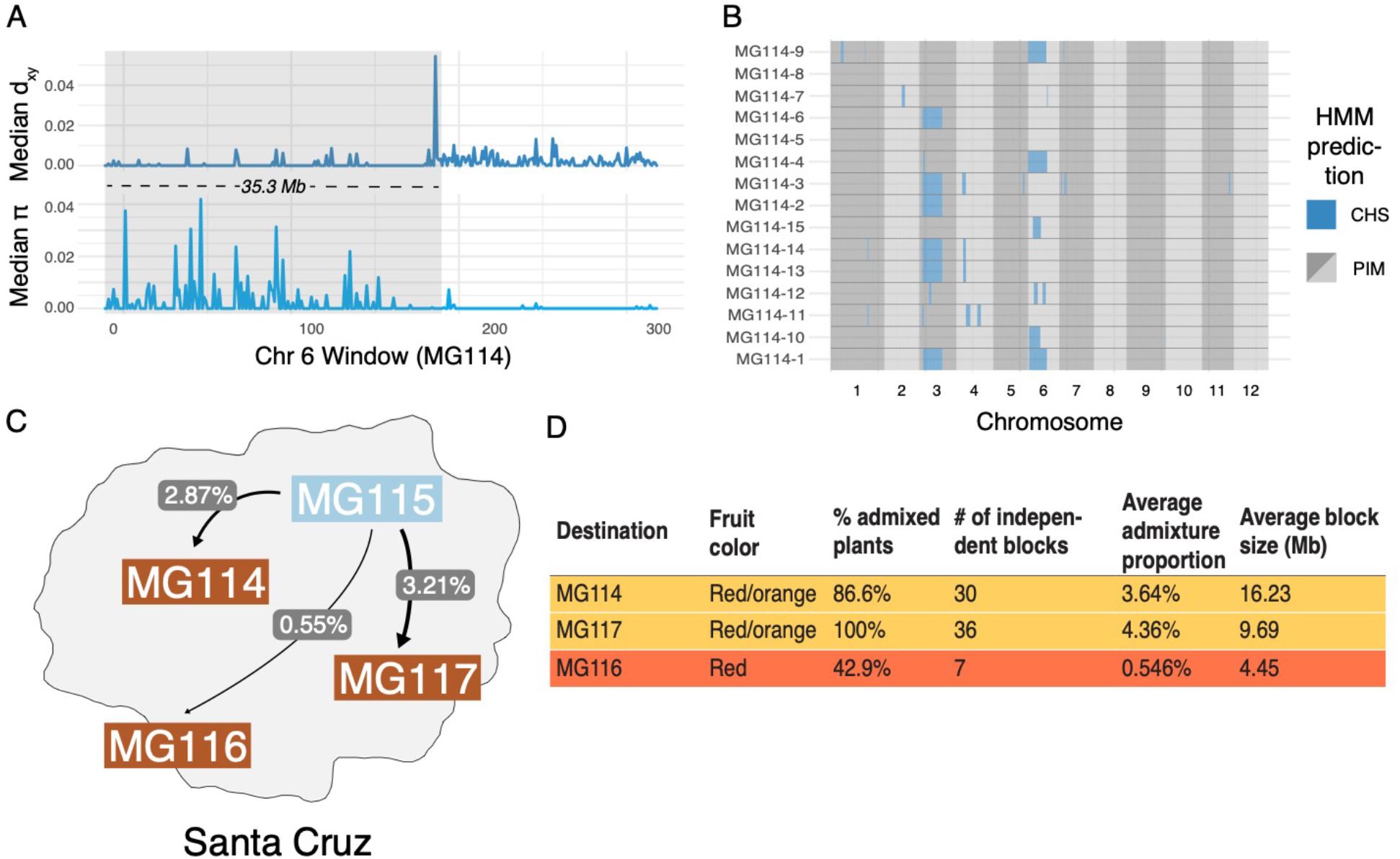
Local ancestry assignment using HMM characterizes a history of endemic x invasive introgression. (A) Patterns of diversity and divergence along chromosome 6 for an MG114 individual. The region of recent coalescence (low divergence; high diversity) with CHS is annotated in gray. This 20.2 kb block segregates at 20% in MG114. (B) Genome-wide HMM predictions for all individuals in MG114. The x-axis is ordered by chromosome and y-axis is ordered by individual. Two large CHS haplotypes segregate at high frequency on chromosomes 3 (40%) and 6 (20%). (C) Visual summary of admixture proportions from CHS into three PIM populations. (D) Summary of HMM assignment for each PIM population. Populations displaying variation in fruit color (MG114 & MG117) have more CHS ancestry than those which are fixed for the ancestral red state (MG116).

**Figure 5:**
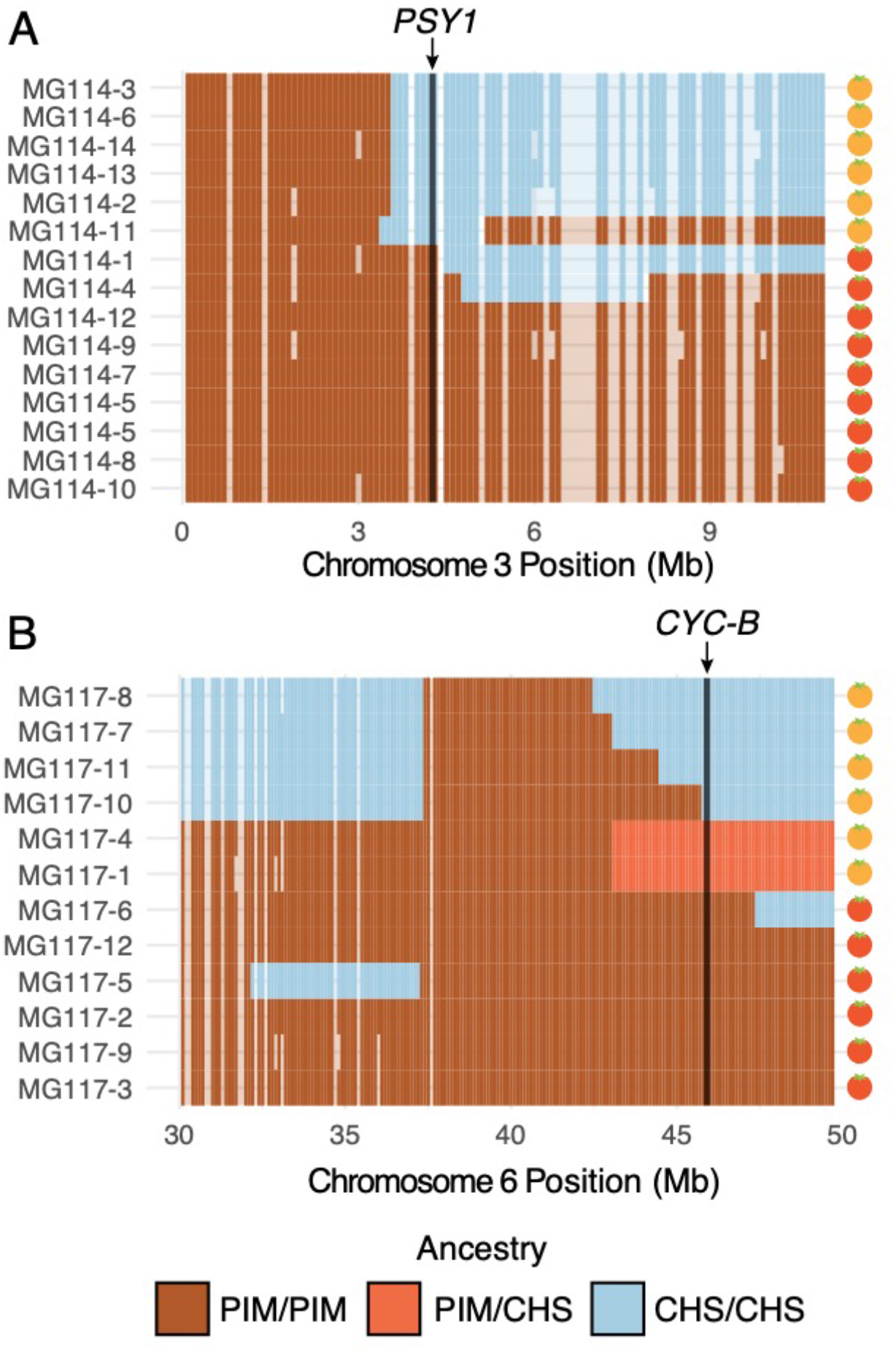
Patterns of local ancestry across focal chromosome regions of MG114 and MG117, enlarged to show variation in introgression block break points at color pathway genes. (A) CHS ancestry at carotenoid biosynthesis gene *PSY1* on chromosome 3 correlates with observed fruit color variation in MG114. (B) CHS ancestry at carotenoid biosynthesis gene *CYC-B* on chromosome 6 correlates with fruit color variation in MG117. Each cell represents 100kb. Empty cells indicate windows with no sequence data. Empty cells are ghost shaded with each ancestry color based on neighboring assignments.

The size of detected ancestry blocks contains information regarding the timing of gene flow, because it depends on the number of recombination events (generations) that have occurred since an initial hybridization event. We can broadly estimate the age of these haplotypes using a simple logarithmic relationship (*SI Appendix, section S5*; Lynch & Walsh, 1998). In MG114 and MG117, age estimates are within the range of 4-12 generations (e.g., the large chromosome 3 and 6 CHS haplotypes in MG114 are estimated at 4.23 and 4.74 generations, respectively). In addition to placing boundaries on when the initial hybridization took place in the past (>4 and up to 12 generations), there is close agreement between age estimates in MG114 and MG117, suggesting that these instances of CHS introgression might have been derived from the same admixture event (*SI Appendix, section S5*).

Relative to the patterns of gene flow from CHS into polymorphic PIM described above, gene flow in the opposite direction (PIM → CHS) was more restricted. The average proportion of PIM ancestry within CHS individuals (at MG115) was 1.62% (s.d. = 2.72%; *SI Appendix, Table S13*). These results point to a potential bias in the direction of gene flow, with more exchange occurring from CHS into PIM than from PIM into CHS.

In addition to inferred introgression between PIM and CHS, we also found evidence for hybridization/introgression involving the other two taxa: GAL and LYC. In particular, we uncovered a recent history of hybridization between CHS and GAL on Isabela—at *la Laguna de Manzanilla* (**Figure 3B & 3D**). These two species have been reported as co-occurring at this site since 2000, and hybridization has previously been hypothesized based on allozyme and morphological analyses (Darwin, 2009; Gibson et al., 2020); **Figure 3D**; *inset images*). Our *fastStructure* and PCA analyses of individuals at this site clearly identify 9 samples as admixed (**Figure 3B**), mostly corresponding with morphological classifications of intermediacy (**Figure 3D**). Of the 9 admixed plants, 4 appear to be first generation backcrosses, whereas 4 are F_2_ (*SI Appendix, Table S7*). We also identified putative cases of CHS/GAL x LYC admixture in two populations on Isabela (**Figure 3B**), although some of these signals might be the product of unmodeled genetic substructure (*SI Appendix, section S3*). On Santa Cruz, low levels of LYC (domesticated tomato) ancestry were also detected within some PIM populations, including a potential hybrid PIM x LYC population MG118 that was predicted to be entirely first-generation hybrids.

### An introgressed origin for orange fruits in PIM

Because introgression from CHS into PIM was most evident in PIM populations that were polymorphic for fruit color (MG114 and MG117), we took an admixture mapping approach to investigate whether introgression influences fruit color variation in these populations. Specifically, we examined the association between local ancestry across the genome and observed fruit color phenotypes, paying special attention to genomic locations of 8 known genes involved in carotenoid biosynthesis (*SI Appendix, Table S14, S15*; Paran & van der Knaap, 2007). For MG117, we found that the presence of orange fruits correlated perfectly with CHS ancestry at only one gene: *CYC-B* on chromosome 6 (**Figure 5B**; *SI Appendix, Table S15*). This association is significant based on a χ^2^ test of independence (χ^2^ = 8.33; df = 1; P = 0.0039). *CYC-B* is a lycopene beta cyclase and the specific locus known to underlie the lighter (orange) colored fruits observed in the endemic species (Stommel & Haynes, 1994).

In contrast, the similarly-sized chromosome 6 haplotype in MG114 (**Figure 4B**) does not include the CHS allele at the *CYC-B* locus. Instead, in MG114 we find that the presence of orange fruits was solely predicted by CHS ancestry at *PSY1* on chromosome 3, the first enzyme in the carotenoid fruit color pathway (*SI Appendix, Table S14*; χ^2^ = 11.12; df = 1; P = 0.0009). Although the role of *PSY1* in the coloration of endemic fruits has not yet been studied, loss of function mutations in *PSY1* have been described in LYC and these produce orange fruit color (Fray and Grierson, 1993). To further investigate the association of *PSY1* with endemic fruit pigmentation, we used previously published RNAseq data (Pease et al. 2016) to examine variation in *PSY1* across the entire wild tomato clade. *PSY1* is expressed at detectable levels in both endemic species and has a highly conserved coding sequence (1203/1239 shared sites), whose exceptions include a single non-synonymous substitution (62R → W) unique to the endemic clade (*SI Appendix, Fig. S14*). This substitution lies outside of the trans-isoprenyl diphosphate synthase protein domain associated with the enzyme’s function, but within the transit peptide signal sequence (residues 1-70), although the specific functional importance of this variant remains to be assessed. Regardless, based on our observed associations between carotenoid biosynthesis loci and fruit color variation, our data support two separate mechanisms underlying the emergence of orange fruits in Galápagos PIM, both of which were likely derived via introgression from CHS.

## Discussion

Biological invasions are one of the foremost threats to global biodiversity, yet we still have a poor understanding of the processes that contribute to invasive colonization success. Here, we studied patterns of genome-wide ancestry and relatedness between endemic, invasive, and continental wild tomato populations in order to reconstruct the history and consequences of a recent biological invasion on the Galápagos. Invasive populations of *S. pimpinellifolium* (PIM) had low levels of genetic diversity and an excess of rare alleles, and we inferred 2-3 recent introductions onto the archipelago from Ecuador and Peru. As a consequence of this invasion, we uncovered evidence for recent and ongoing gene flow between PIM and the congeneric endemic species *S. cheesmaniae* (CHS). Local ancestry at two key carotenoid loci further supported an introgressed (CHS) origin for orange fruits in at least two invasive populations.

Together, our results reconstruct the history of invasion, and infer that the source of convergent phenotypic evolution in the invasive populations is introgression of important functional alleles from endemic relatives.

### Genomic data reconstruct the demographic history of invasion onto the Galápagos Islands

Our analyses identify 3 independent introduction events, yet only a single event from Ecuador comprises 98% of sampled invasives on the archipelago. The other two introductions—each represented by single plant collections—either did not produce large invasive populations or they result from much more recent introductions which have not yet established broadly. Indeed, our PCA (**Figure 2A**) supports the idea that they are the product of more recent introduction events, as these two collections are more closely related to the mainland PIM populations than plants derived from the primary introduction.

For the primary introduction, although we observed variance in source region predictions (*SI Appendix, Fig. S2*), the consensus prediction supports a southcentral origin near Guayas or Los Rios provinces. Intriguingly, historical data on human migration and trade on the islands also point to this as a likely region for the source of invasive PIM. First, although Ecuadorian colonizers of the Galápagos originate from across the country, one of the earliest and largest bursts of migration coincided with the Tungurahua province earthquake in 1949 (Toral-Granda et al., 2017). This province is geographically central and close to Los Rios. Second, the vast majority of all trade between Galápagos and the continent occurred—and continues to occur—from Guayaquil, the second largest city in Ecuador (Lundh, 2004; Toral-Granda et al., 2017). The surrounding agriculture regions, which includes Los Rios, would be the most proximate sources for raw product shipments to the islands. This historical context provides additional support for our genetic inference of a majority southcentral origin for invasive populations.

Our inferences also clearly implicate humans as the source of PIM introduction. Our demographic reconstruction points to a recent bottleneck and expansion of PIM on the archipelago (**Table 2**), much more recent than our estimate for CHS. Similarly, our inference that LA0411 (a mainland Ecuador accession) is the product of back migration from the Galápagos underscores the recent and likely substantial human influence on the movement of PIM. We conclude that PIM is most likely the result of a recent, human-mediated expansion on the archipelago. Human introduced species represent upwards of 70% of all alien plant species on the Galápagos (Quiroga, 2018), and PIM has similarly been hypothesized to be the product of a human introduction; however the timing and mode of its introduction—including the role of humans—was not previously known.

### Hybridization as a consequence of invasion onto the Galápagos

One key evolutionary consequence of PIM’s introduction onto the Galápagos that emerges from our analyses is its hybridization with endemic congeneric species— primarily CHS. Hybridization has been hypothesized as a mechanism for promoting invasive colonization success, as it could help overcome the adaptive limits that might otherwise be imposed by genetic bottlenecks during the colonization process [the so-called “genetic paradox” of invasion (Allendorf & Lundquist, 2003; Sakai et al., 2001)). These bottlenecks can be especially severe during introductions onto islands (Kolbe et al., 2004; Colautti et al., 2005; Golani et al., 2007). In addition, several factors indicate the high potential for gene flow specifically between the four studied species (CHS, GAL, PIM, LYC), including their very close evolutionary relationships (all are members of the red-fruited Esculentum subclade within the wild tomatoes; Pease et al., 2016), and their incomplete reproductive barriers (Rick, 1956; Rick & Bowman, 1961; Rick & Fobes, 1975). Nonetheless, previous analyses based on handfuls of loci provided conflicting evidence for and against the occurrence of gene flow between species presents on the island (Nuez et al., 2004; Darwin, 2009).

Our data provide clear evidence for recent hybridization and introgression between all four tomato taxa on the archipelago. Although our focus here is primarily on CHS and PIM, we also find evidence for recent hybridization and/or introgression between CHS and PIM (Santa Cruz), PIM and LYC (Santa Cruz), CHS and GAL (Isabela), and, to a lesser extent, CHS and LYC (Isabela). These patterns suggest that hybridization—both with congeneric endemics (CHS and GAL) and invasives (LYC)— could serve as a source of adaptive genetic variation in invasive PIM.

The most prominent signal of gene flow is between PIM and CHS (**Figure 3; Figure 4; Figure 5**), including clear evidence for both early generation (F1 and F2) hybrid offspring as well as older introgression 4-12 generations in the past. Our results indicate (i) that CHS ancestry is maintained in some PIM populations beyond initial hybridization and (ii) that gene flow is ongoing.

The potential consequences of secondary genetic contact are numerous (Wolf et al., 2001; Todesco et al., 2016). While we do not have direct data on relative fitness of hybrids, the persistence of later generation CHSxPIM hybrids indicates they are not immediately selected against. Indeed, the genomes of most admixed PIM (MG114 and MG117) are consistent with a history of secondary contact and gene flow characterized not by strong hybrid incompatibility, but a less restricted exchange of alleles between species. Furthermore, the nonrandom distribution of CHS ancestry throughout admixed PIM suggests that it may be selectively maintained in certain regions of the genome.

Instead of observing a heterogeneous set of CHS alleles in the backcrossed genome of PIM, we find that CHS ancestry is enriched on chromosomes 3 and 6, and absent in much of the rest of the genome, in both MG114 and MG117 (**Figure 4B**; *SI Appendix, Fig. S12*). Moreover, our local ancestry predictions provide evidence that these introgressed regions may contain key genes responsible for the emergence of orange fruit color in MG114 and MG117.

### Orange fruit color in island PIM was derived via introgression from CHS

A key finding of our analyses is that introgression is likely the source of phenotypic convergence on orange fruits that is observed in invasive Santa Cruz PIM. Orange/yellow fruit color is diagnostic for the endemic species (CHS is typically pale yellow; GAL is typically orange) but extremely rare in PIM. At the genetic level, convergence could be based on three potential sources of variation: ancestrally segregating variation, introgression, or via a *de novo* transition. Of these, ancestral variation is the least likely: The very few described examples of orange fruits among continental PIM are all located in Peru (e.g. Sifres et al. 2007), and none have been reported in the inferred geographic region of origin of this invasion (TGRC passport data; www.tgrc.ucdavis.edu). One goal here was therefore to distinguish between introgressed and novel mutation as the source of phenotypic convergence. We did so by mapping the landscape of introgression throughout the genomes of invasive PIM plants, and evaluating its association with observed fruit color variation, and with loci known to underlie this trait in Solanum (Paran & van der Knaap, 2007). With these data we inferred a unique scenario in which phenotypic transitions to orange fruits in two different invasive PIM populations were each derived from introgression at a distinct carotenoid locus: *CYC-B* or *PSY1*.

Our data in conjunction with existing experimental evidence indicate that *CYC-B* is the causative locus for orange fruits in MG117. Interestingly, *CYC-B* mutants were first identified as natural allelic variation in the endemic species CHS (Rick, 1956).

Introgression of the CHS *beta* allele at *CYC-B* into LYC causes the accumulation of β-carotene in ripening tissues and the production of orange fruits (Stommel & Haynes, 1994). Orange fruits segregate as a single dominant gene, and genotypic variation at this locus explains a large fraction of fruit color variation in experimental crosses (Rick, 1956; Stommel & Haynes, 1994). Our data show a clear association between CHS ancestry and orange fruit color at this locus (**Figure 5**)—including the observation that individuals heterozygous for ancestry display the dominant phenotype—so we infer that introgression of CHS *CYC-B* into PIM has the same large, dominant effect on fruit color in admixed individuals of this wild species.

Unlike *CYC-B, PSY1* was first identified in the spontaneous fruit color mutant *yellow-flesh* in LYC (accession LA2997; Fray and Grierson, 1993), and its role has not been directly evaluated in CHS or GAL. Recessive *r* mutants at *yellow-flesh* carry a truncated version of *PSY1* that is unable to convert precursor into phytoene. The resulting fruits accumulate almost no carotenoids and the yellow skin pigmentation is driven primarily by the accumulation of the flavonoid chaloconaringenin (Fray and Grierson, 1993). Using previously published RNA-seq data (Pease et al., 2016), we confirmed the expression of *PSY1* in both endemic species and did not detect any truncation or premature stop mutations. Rather, we identified a single non-synonymous substitution (62R→W) within the transit peptide signal domain, found in both endemic species. Disruption of the transit signal sequence may prevent localization to the chloroplast and thus result in a nonfunctional enzyme, although determining the exact role *PSY1* has in endemic—and by extension PIM—fruit coloration would require future functional confirmation. Regardless, *CYC-B* and *PSY1* in invasive orange-fruited PIM are unequivocally derived from CHS, and current functional knowledge of both loci indicate their effects on fruit color could entirely explain observed phenotypic variation in orange-fruited PIM.

Finally, the ubiquitous lighter (orange and yellow) fruits of the two endemic species, the appearance of convergence towards endemic-like fruit colors in invasive PIM, and the likely independent recruitment of endemic fruit color alleles at *PSY1* and *CYC-B* in MG114 and MG117, together suggest intriguing evidence that lighter fruits may have a specific selective advantage on the islands. The potential environmental basis of this selection is unknown, however differences in fruit dispersal—including disperser color preference(s) and/or fruit color apparency—on the islands vs the continental mainland, could be a likely mechanism. Alternatively, at least in the case of *PSY1* which likely involves either a full or partial loss of function, orange pigmentation could arise due to relaxed selection, if it is more costly to produce red fruits and they have no specific advantage in island environments. Future field experiments and fitness measurements will help to distinguish among these selective hypotheses.

### Conclusions

Overall, our results reconstruct a complex and recent history of invasion by wild tomato onto the Galápagos Islands, and highlight the potential importance of gene flow during colonization. Our results also add to an emerging phenotypic convergence literature by highlighting how admixture brought on by anthropogenic change can drive convergence over very short time scales. While the adaptive benefit of orange fruits remains to be evaluated, our finding of two separate molecular mechanisms underlying orange coloration each derived from CHS is highly suggestive that lighter fruit pigmentation is favored in the island environment. This study underscores how the long history of research on the Galápagos Islands continues to enrich our understanding of evolutionary processes in the natural world.

## Materials and methods

### Population sampling and genotyping

We sampled leaf tissue from 13 wild populations of invasive and endemic tomato taxa on the three largest islands of the Galápagos archipelago: San Cristobal, Santa Cruz, and Isabela (**Figure 1**; *SI Appendix, Table S1*). Leaf tissue was dried in silica and DNA was extracted using Qiagen Plant Mini Kits (Qiagen, Valencia, Calif., USA). Two double-digest restriction site associated DNA sequencing (ddRAD) libraries were prepared using *PstI* and *EcoRI* enzymes by the Indiana University Center for Genomics and Bioinformatics. Libraries were sequenced across two Illumina NextSeq flowcells (150 bp, paired-end, mind-output).

Raw reads were filtered for quality, trimmed of adapter sequence and low-quality bases using *fastp* (Chen et al., 2018), and demultiplexed by individual using the *process_radtags* program in *Stacks* (ver. 2; Catchen et al., 2013). Reads were mapped to the *S. lycopersicum* reference genome version SL3.0 using BWA (Li & Durbin, 2009). Bam files of 140 continental accessions representing the full species range of PIM (*SI Appendix, Fig. S1 and Table S2*) were jointly reanalyzed with the new samples in *Stacks*. Mapped reads were assembled and variants were called with the Stacks *ref_map* pipeline. Genotype calls made with fewer than 8 reads were removed and subsequently we retained only sites having data for at least 80% of all 306 individuals. For all analyses except diversity/divergence calculations (π, Tajima’s D, d_XY,_ F_ST_) and *Treemix*, we pruned sites in high LD (r^2^ > 0.7) using *bcftools*. All scripts are available at http://github.com/gibsonmatt/galtom.

### Nucleotide diversity and divergence estimates

Within-population diversity and divergence estimates across the genome were calculated using the *Stacks* program *populations*. Windowed π (**Figure 1C)** was extracted from *Stacks* output. For pairwise comparisons between the islands, Ecuador, and Peru (right panel **Figure 1C**), we calculate genome-wide pairwise divergence directly from the assembled RAD loci (samples.fa file) using a custom Python script (http://github.com/gibsonmatt/galtom). For each pairwise comparison between samples, we count the total number of sequence differences and total number of sites for which both samples have data. This choice allows us to conveniently model patterns of diversity between diploid samples in our introgression HMM using a binomial. We also calculated the average genetic distance from each accession to all Galápagos PIM across polymorphic sites in the R package *adegenet* (Jombart & Ahmed, 2008; **Figure 1A**).

### Phylogenetic reconstruction

We inferred a maximum likelihood tree of population relationships (**Figure 1B**) using *Treemix* with no specified migration. The *Treemix* input file was generated from a VCF using a custom Python script (http://github.com/gibsonmatt/galtom). We expected abundant phylogenetic discordance both within Galápagos PIM (given its recent divergence from Ecuador) and within the red-fruited tomato clade in general (Pease et al., 2016). To this end, we generated 1,000 replicate datasets by sampling (with replacement) 500 SNPs from the full dataset. These trees were visualized using *densitree* as implemented in the R package *phangorn* (Schliep, 2011). Both the set of replicate trees and the consensus tree were rate smoothed using *r8s* (λ = 1), using the chronopl function available in the R package *ape* (Paradis & Schliep, 2019). In addition to *Treemix*, we also inferred a maximum likelihood phylogeny of individual sample relationships using *RAxML* (*SI Appendix, Fig. S3*). We subset our dataset to one individual per population for Galápagos collections and to 15-20 samples per geographic region (Peru and Ecuador) for mainland accessions. We ran RAxML using the GTRCAT approximation of the general time reversible model of substitution allowing for rate heterogeneity. 25 alternative runs from distinct maximum parsimony trees were performed, from which we selected the best single tree.

### Demographic inference

We modeled bottleneck demographic histories from the site frequency spectrum (SFS) using *δaδi* (Gutenkunst et al., 2009), calculating the folded SFS using *easySFS* (https://github.com/isaacovercast/easySFS). For population MG114, we down sampled from 30 to 14 chromosomes, striking a balance between the number of segregating sites and levels of missing data in frequency bins. We chose this population because the larger number of samples (2N = 15) relative to other PIM populations afforded more statistical power. Prior to calculating the SFS for MG114, we removed regions inferred as introgressed by our HMM, as we found that a large fraction of singleton and doubleton sites in MG114 were shared with CHS (*SI Appendix, section S6*). These sites would be spuriously interpreted as *de novo* mutations derived in PIM post-colonization, thereby biasing our parameter estimates. See *SI Appendix, section S6* for further discussion of our filtering scheme and its effects on *δaδi* estimates. For population MG115, we down sampled to 24 chromosomes. For each population, two models were evaluated: (1) a neutral model of no population size change and (2) an instantaneous bottleneck model. The parameters of the bottleneck model are described in the results. We use an extensive three-step optimization pipeline described in Portik et al. (2017) to explore the demographic parameter space of the PIM bottleneck model. For each successive round, we increase the number of replicate runs (10, 200, and 500) and decrease the amount of parameter perturbation (3, 2, and 1-fold). The parameters from the replicate with the highest likelihood in each round are taken as the starting parameters in the next round. To estimate confidence intervals for our parameters, we performed a nonparametric bootstrapping procedure (assuming independence among sites) by sampling randomly from the observed SFS 2,000 times. Since we filter for LD, we consider this method to be appropriate.

Time in *δaδi* is represented in units of 2N_ref_ generations. To convert from these coalescent time values to numbers of generations, we estimate N_ref_ as θ/4μL, where θ is the population scaled mutation parameter (estimated by *δaδi*; 99.95 for MG114), μ is the per-generation mutation rate (assumed here to be 1⨯10^−8^ muts/bp/generation), and L is the length of queried sequence (5,806,952 bp; estimated from the data as the total number of bases where at least one sample in the focal population had data).

Bottleneck and final effective population sizes were similarly converted from relative to absolute values using N_ref_.

### Inferring gene flow between contemporary island populations

We used several independent methods to characterize genetic structure in our dataset. First, we applied *Treemix* (Pickrell & Pritchard, 2012) to assess evidence for broad signatures of admixture among populations. Since we were interested in understanding the history of PIM, we subset our full SNP dataset to exclude LYC populations since the exact taxonomic status of these samples are unclear and do not help address questions of gene flow between PIM and endemic species. Furthermore, we remove PIM populations with fewer than 8 samples. We ran *Treemix* with several values for *m* (0-8), the migration parameter which determines how many reticulation branches are allowed. Based on the likelihoods provided by *Treemix*, we determined that a migration parameter of 6 was most appropriate. Increasing *m* led us to infer more intraspecific migration within PIM, but no additional interspecific events could be inferred and the increase in data likelihood was marginal (*SI Appendix, Fig. S7 and Table S9*).

We also employed model-based (*fastStructure* ([Raj et al., 2014]) and non-model-based (multi-locus PCA) methods to evaluate genetic structure. We ran both methods using the LD-filtered dataset of 5,767 SNPs, and on each island separately. For each island, we ran *fastStructure* for values of *k* between 1 and 7. We then chose an appropriate model complexity using the *chooseK*.*py* script supplied with *fastStructure*. After a value for *k* was chosen, we evaluated the stability of the ancestry assignments across 5-10 separate runs. The multi-locus PCA was implemented in the R package *adegenet*.

For select populations with evidence for admixture, we ran *NewHybrids* (Anderson & Thompson, 2002) to identify any early generation hybrid individuals (F_1_, F_2_, and backcrosses). *NewHybrids* was ran using the same LD-filtered dataset as in *fastStructure* and PCA (script for converting from vcf to *NewHybrids* format is provided at http://github.com/gibsonmatt/galtom). For each *NewHybrids* run, we specify three groups: a parent population A group, a parent population B group, and an admixed group. For the CHSxPIM interaction on Santa Cruz, parent populations were defined as non-admixed MG115 and MG113 individuals. For the CHSxGAL interaction on Isabela, parent populations were MG120C and MG120G. For each population we ran the Markov chain for at least 6,000 iterations. Individual assignments were not sensitive to the choice of priors. Lastly, four-sample D-statistics (Durand et al. 2011) were calculated using the *compD* software package (https://github.com/stevemussmann/Comp-D), using *S. pennellii* (LA3778) as an outgroup species.

### Introgression analysis

We implemented a hidden Markov model (HMM) to identify fine-scale genomic signatures of introgression. Although RAD sequencing is not optimal for genome scanning due to lower marker density, recent signatures of introgression should be large and detectable. Nonetheless, we acknowledge the fact that our sequencing methods may not allow for full characterization of the landscape of introgression. In our analysis we leveraged the fact that, in regions of recent introgression, genetic diversity within the destination population (π) should be elevated relative to diversity between the source and destination populations (d_XY_). In other words, regions of recent interspecific coalescence should resemble the introgressing species more than the destination species. We calculate these divergence/diversity estimates directly from assembled RAD loci (samples.fa file from Stacks).

Our HMM featured three hidden states: (A) CHS ancestry, (B) PIM ancestry, and (C) heterozygous CHS/PIM ancestry, for which we used π, d_XY_, and the mean of π and d_XY_ (*m*) in nonoverlapping 100kb windows, respectively, to calculate emission probabilities. We used three binomial models to obtain these probabilities (see *SI Appendix, section S4* for a detailed description of the model). For each chromosome and for each focal comparison, we found the most likely hidden state path for a given sequence of π and d_XY_ using the Viterbi algorithm, controlling for underflow by operating in log-space. Because of the coarse scale (100kb windows) used in ancestry assignment, our HMM struggled to identify smaller genetic signals potentially consistent with introgression. *SI Appendix, Fig. S9* provides a per-window look at how ancestry assignment correlated with patterns of diversity and divergence in MG114. Patterns of diversity/divergence vary substantially even between adjacent windows and thus it is likely that we are not capturing sub-100kb signals of introgression. Nonetheless, the large size of the blocks we do identify allows us to be confident they are the result of introgression rather than ILS.

## Acknowledgements

We thank the Galápagos Science Center staff on San Cristóbal for logistic and permitting support, and the Galápagos National Park for assistance locating and sampling endemic populations. On-site field support was provided by Marcelo Loyola and Genaro Garcia. This work was supported by a United States National Science Foundation award (IOS 1127059) to LCM and the Indiana University Brackenridge award to MJSG. All field collections were made with appropriate permits and prior authorization by the Galápagos National Park and Ecuadorian Ministry of Environment (Permits PC-40-18 and PC-72-19).

