## Supplementary material for "Reconstructing the history and biological consequences of a plant invasion on the Galápagos islands": SI Appendix

PNAS
www.pnas.org

Supplementary Information for

### Reconstructing the history and biological consequences of a plant invasion on the Galápagos islands

Matthew J.S. Gibson, María de Lourdes Torres, Yaniv Brandvain, & Leonie C. Moyle

Corresponding Author: Matthew JS Gibson


**This PDF file includes:**

Supplementary text
Figures S1 to S14
Tables S1 to S16
SI References

**Supplementary Text**

**S1. Field collections and taxonomic treatments**

**S1.1. Collections and DNA extraction.** We visited San Cristóbal, Santa Cruz, and Isabela—the three most populated islands in the archipelago. These islands contain 40 documented TGRC locations (TGRC passport data) and these sites, as well as those described in Darwin (2009) and Nuez et al. (2004), were searched during our expeditions. We looked for plants within a 200m radius of each previously documented collection site, and we also searched for additional undocumented populations. Populations were sampled across linear transects and detailed photographs of leaf, fruit, and flower morphology were taken of all sampled individuals against color and length standards. The latitude and longitude of each collection location was logged using a smartphone. Each site and the identity of species found within are described in **Table S1.** For extracting DNA, 3-5 leaves were sampled from each plant and immediately dried in silica gel. Tissue disruption was performed with mini-pestles and DNA for each sample was extracted using Qiagen Plant Mini kits (Qiagen, Valencia, Calif., USA) at the Galápagos Science Center (San Cristobal, Galápagos, Ecuador). Our sequencing pipeline is described in the main text.

**S1.2. Taxonomy.** Population identity was determined based on the taxonomic treatments described in Darwin et al. (2003), with the exception of LYC, which we separated into two forms—LYC (domesticated tomato) and LYC var. *cerasiforme* (CER; cherry tomato; Rick, 1956)—to maintain consistency with Nuez et al (2004). Ripe fruit color was the primary trait used in species identification, with the exception of individuals designated as orange-fruited *S. pimpinellifolium*, based on all other characters. We additionally measured nine leaf traits [leaf length, leaf width, terminal leaflet length, leaftlet count, interjected leaflet count, petiole length, internode length, leaflet length, and leaflet width previously identified as diagnostic by Darwin (2003).

**S2. LA0411 and our inference of back migration**

**S2.1. Original collection notes for LA0411.** Charles Rick, 1956: “Pichilingue. 200 m. L. pimpinellifolium or L. esculentum var. cerasiforme? Weedy vigorous form growing as weed in corn field N of station buildings. Population of 15 plants found here. 5 had ripe fruit. Long internodes, leaves more elaborate than ? pimp. Few long hairs at growing point like Santa Cruz pimpinellifolium. Alternation all = 3 or +/- 3. Flowers small, stigmas all exserted 1 mm or more. Population seems to be entirely uniform. Inflorescence all 1st order, racemose. Fruits large for pimp. = 1.5 cm, red. No vectors seen. Long tailed thick billed black birds seen near patch. These said to eat wild tomatoes near Guayaquil cement factory. Odor = esculentum. Anthocyanin = dark”.

**S2.2. Evidence for back migration.** Based on patterns of genomic relatedness to Galápagos and surrounding PIM accessions, we infer that Ecuadorian accession LA0411 (collected in 1956 by Charles Rick) is the product of a back migration from the Galápagos to mainland Ecuador. This accession showed a particularly strong resemblance to Galápagos PIM (average dxy = 0.0006) and was also divergent from neighboring mainland Los Rios accessions (**Figure S4**). This accession was indeed noted as morphologically similar to Galápagos PIM tomato collections when sampled on mainland Ecuador in 1956 (TGRC passport data; http://tgrc.ucdavis.edu; S2.1, above). This inference has two implications for understanding the history of PIM and other invasive species on the islands. First, it sets an upper bound on the timing of the initial introduction of PIM to the Galápagos as no later than 1956, as it must have already been established there prior to a back migration event. Second, it highlights the substantial connectivity between this region (the putative source of invasive PIM) and the Galápagos in general.

This finding does not affect our analyses of invasive population origins, including our inference that most invasive PIM have an Ecuadorian source. Invasive PIM have high genetic similarity with many accessions in this region of Ecuador (**Figure 2A**). Furthermore, running *Locator* without accession LA0411 produces results identical to those described above.

**S3. Further details on evidence for gene flow**

We use several statistical methods for detecting gene flow between invasive and endemic populations. Our primary focus was on characterizing admixture between CHS and PIM on Santa Cruz (described in text), however several additional patterns are also worth presenting. These are discussed below.

**S3.1. Treemix.** In addition to the two cases of interspecific PIM X CHS admixture, three cases of intraspecific admixture within PIM were inferred using *Treemix*: one from Peru PIM into Ecuador PIM, one from the ancestral Galápagos branch to MG107, and one from Ecuador PIM to MG111. It is difficult to interpret the factors responsible for these inferred events, although they may not all reflect true admixture cases. The edge between Ecuador and Peru PIM is most likely a byproduct of our distinction between these groups; in reality the relatedness among mainland samples reflects a pattern of IBD across latitude. The edges leading to MG107 and MG111 could suggest the occurrence of additional minor introduction events nested within the major Ecuador event. Such a scenario might be plausible given the known substantial trade between the Galápagos and central mainland Ecuador, and the occurrence of at least one reintroduction event. However, we have no independent support for such additional events. Increasing *m* in *Treemix* had the effect of inferring additional intraspecific migration within PIM, but did not infer any additional between species admixture (see **Figure S6** for additional *Treemix* run summaries) and the increase in data likelihood was marginal (**Table S8**). This indicates that the precise parameter choices in these analyses do not change the number of introgression events inferred between invasive PIM and the endemic island species, beyond those that described in the main text.

**S3.2. *S. lycopersicum*.** In addition to MG118 (population of F1 PIM x LYC plants) described in text, other weaker signals of LYC ancestry in PIM (e.g., in MG113, 114, 116, & 117; **Figure 4A**) may also be the result of post-colonization admixture, however they may also represent latent (unmodeled) population structure within PIM and/or reflect the hybrid ancestry of LYC var. *cerasiforme* (Ranc et al., 2008). Increasing K from 3 to 4 in *fastStructure* swapped the minor LYC ancestry fractions seen in MG113, MG116, and MG117 for a fourth ancestry class. However, the minor LYC component seen in MG114 as well as the 50/50 LYCxPIM MG118 population retained their original LYC classifications. It is difficult to interpret this given the complex and unresolved history of LYC var. *cerasiforme* (Ranc et al., 2008), however it may indicate (i) the presence of two or more distinct LYC lineages on the Galápagos or (ii) a contribution by *S. galapagense* (GAL; not sampled on Santa Cruz but historically present). The former scenario may be expected if multiple varieties of domesticated seed have been used for cultivation on the archipelago.

Separate from the clear case of CHS x GAL admixture on Isabela at *la laguna de manzanilla* (discussed in the main text), additional signals of admixture between CHS/GAL (MG120) x LYC (MG122 and MG126) were also detected (**Figure 4B**). In MG122, increasing K from 3 to 4 clearly separates the three inferred GAL x LYC individuals into a separate non-admixed sub cluster, suggesting that the inferred GAL ancestry in these samples likely reflects latent substructure within LYC. In contrast, even when increasing K from 4 to 5, the three inferred CHS x LYC samples in MG126 retain their minority CHS component, consistent with a real signal of shared ancestry between CHS and LYC for these individuals. Whether this is the result of gene flow post-colonization of LYC on the islands or a reflection of breeding history (CHS germplasm has been used as a source for certain beneficial crop traits; Rick, 1967; Stommel and Haynes, 1994) is unclear.

#### S4. Local ancestry assignment with Hidden Markov Models

Our HMM was implemented in R and Python (code available at http://github.com/gibsonmatt/galtom). We used binned pairwise sequence divergence in 100kb nonoverlapping windows to define emission probabilities and defined three hidden states (homozygous CHS ancestry, homozygous PIM ancestry, and heterozygous CHS/PIM). We use our HMM to identify regions of recent coalescence between each population.

For a focal individual and in each window we calculated emission probabilities using three binomial models:

$$P_{CHS} = \binom{d_1}{s} \pi^s (1-\pi)^{d_1 - s}$$

$$P_{PIM} = \binom{d_2}{s} d_{XY}^s (1-d_{XY})^{d_2 - s}$$

$$P_{HET} = \binom{d_3}{s} m^s \, (1-m)^{d_3 - s}$$

where $s$ is the number of sites in the window and $d_1$, $d_2$, and $d_3$ are the median number of differences to MG114, MG115, and the mean of $d_1$ and $d_2$, respectively. Transition probabilities (for the CHS into PIM model) were defined as follows:

$$P[CHS - CHS] = ([1-r] + r) + a$$

$$P[PIM - PIM] = ([1-r] + r) + (1-a)$$

$$P[CHS - PIM] = P[PIM - CHS] = P[HET - CHS] = P[HET - PIM] = r \times a$$

where *r* and *a* can be interpreted as proportional to the per-window recombination rate and admixture proportion, respectively. We scaled *r* by a factor *t*, which can be interpreted as proportional to the time since admixture. Increasing *t* will cause the HMM to be more likely to switch between states since recombination will have had more time to break up introgressed blocks. We found that our HMM is relatively insensitive to chosen transition probabilities. For all populations we chose to use 0.002, 0.48, and 10 for *r, a,* and *t*, respectively, as these produced consistent annotations that agreed with patterns of observed nucleotide diversity. Each focal individual in a population was analyzed separately against each of the individuals in the potential donor population. For example, to analyze introgression from CHS → PIM, divergence to all MG114 and MG115 individuals is calculated and used to define emissions.

**S5. On the timing of introgression**

The size of introgression blocks contains information about their age. Over time, recombination will break up contiguous blocks into smaller tracts, resulting in a mosaic of ancestry throughout the genome. We can roughly estimate the age of any given block if we assume that an initial hybridization event was followed by subsequent backcrossing to non-admixed parental individuals. Using a simple logarithmic relationship, the expected time *t* required to arrive at a proportion *p* of the donor genome after repeated backcrossing is estimated as:

$$\log\left(\frac{1}{p}\right)/\log(2)$$

via Lynch & Walsh (1998). As block size gets smaller, the expected time since hybridization increases.

The distribution of introgression tract sizes and the concordance of break points between blocks within and between populations is indicative of non-independence among blocks. Such a pattern makes any detailed assessment of the timing of these events difficult, and our broad estimate makes several simplifying assumptions. The occurrence of selfing and/or small population sizes may also affect the relationship between block length and age of introgression. Nonetheless, it is clear that many of the observed patterns of CHS ancestry throughout the genomes of MG114 and MG117 plants were derived from very recent (and likely shared) events. We further investigated the degree to which these histories of introgression were shared by comparing the ancestry of each population window-by-window. Specifically, we called windows of shared CHS ancestry in MG114 and MG117 as those where at least one individual in each population was assigned as CHS by our HMM. Across the genome, 250 windows were inferred to be of CHS ancestry in both populations, representing 26.8% of all CHS ancestry in MG114 and 35.4% of all CHS ancestry in MG117. These patterns imply a complex and partially shared history of admixture in MG114 and MG117. This inference is consistent with several of our other analyses, including *Treemix* (**Figure 4**) and D-statistics which also pointed to a shared basis to detected patterns of admixture. At the resolution which we are able to assign local ancestry, our data also firmly indicate that endemic variation is maintained within PIM beyond the first or second generation of hybridization.

**S6. The impact of recent introgression on demographic inferences with *δaδi***

A recent history of introgression from CHS/GAL into invasive PIM populations could bias *δaδi* parameter estimates towards older dates by introducing low frequency alleles that would incorrectly be interpreted as *de novo* mutations derived post-expansion. To examine this possibility, we determined whether rare alleles in MG114 were private (i.e., informative of the time since bottleneck) or shared with CHS (e.g., from introgression). We found that a large portion (52%) of singleton and doubleton SNPs in MG114 were shared with CHS (MG115), suggesting that introgressed variants could substantially affect our parameter estimates of the occurrence and timing of demographic changes within invasive PIM. For this reason, prior to model fitting we removed SNPs within all genomic regions where at least one MG114 individual had evidence for CHS ancestry based on our HMM. Any region inferred to be of CHS ancestry in any individual was removed from the dataset for all individuals. This resulted in 365 sites being filtered from the SFS relative to the original dataset. The results of model fitting using the masked data are reported in the main text while the unmasked dataset results are reported in **Table S16**.

As expected, θ was higher in the unmasked dataset including introgression (108.094) than in the masked dataset (99.95). Accordingly, Nref was also higher in the unmasked dataset (465.36) compared to the masked (430.32). The total time since the bottleneck was inferred to be substantially older using the unmasked SFS (a difference of 1894.45 generations). NF (recovery population size) and F (inbreeding) were also heavily affected by filtering of introgression regions. The optimal NF and F were 335.06 and 0.0008, respectively, in the unmasked dataset and 2730.97 and 0.24, respectively in the masked dataset. These results confirmed our expectation, based on allele sharing, that including introgressed genomic locations substantially inflated *δaδi* parameter estimates.

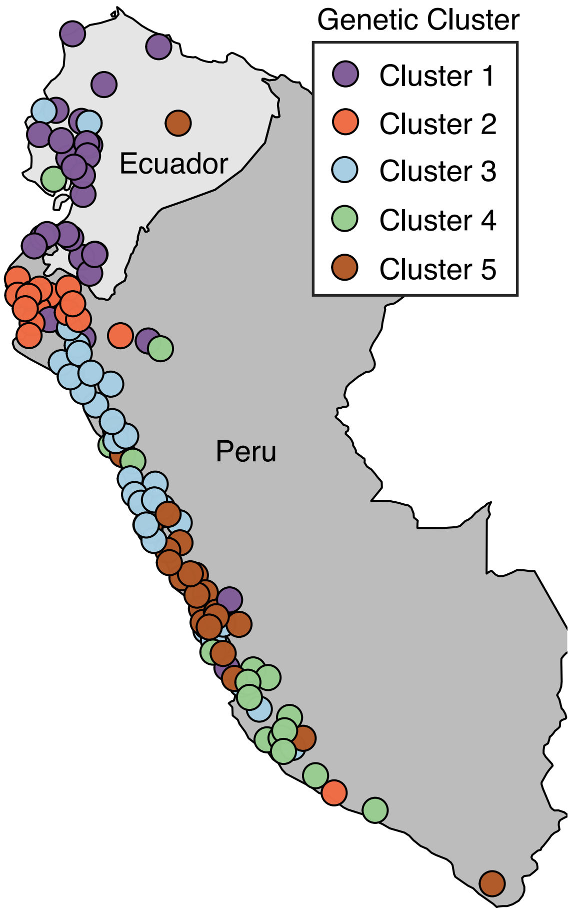


**Figure S1:** Map of mainland collection sites, colored by genetic ancestry cluster as determined in Gibson & Moyle (2020).

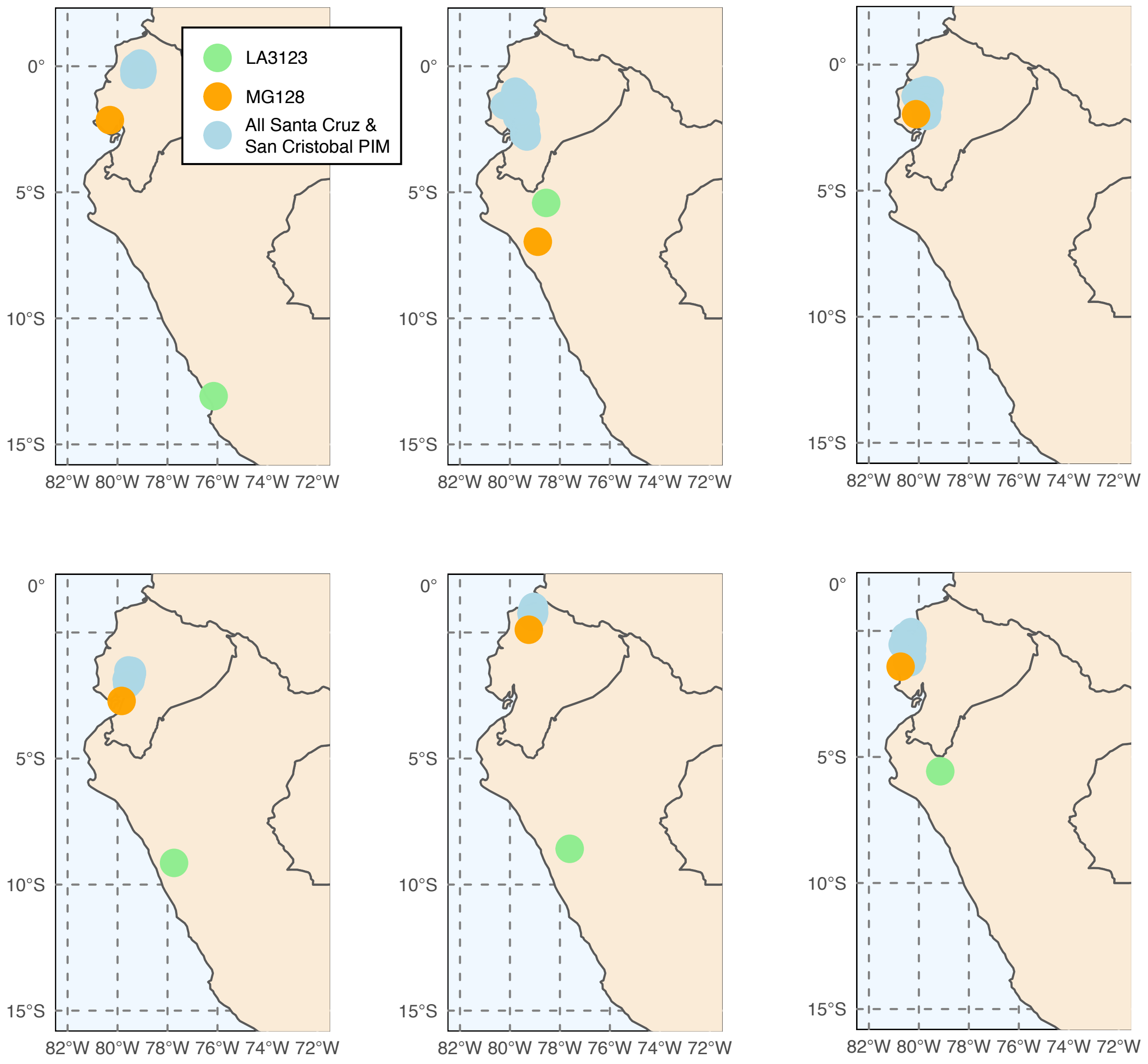


**Figure S2**: Six runs of *Locator* (Battey et al., 2019) generally support a three-invasion scenario. Exact source localities for MG128-1 and LA3123 varied substantially across run.

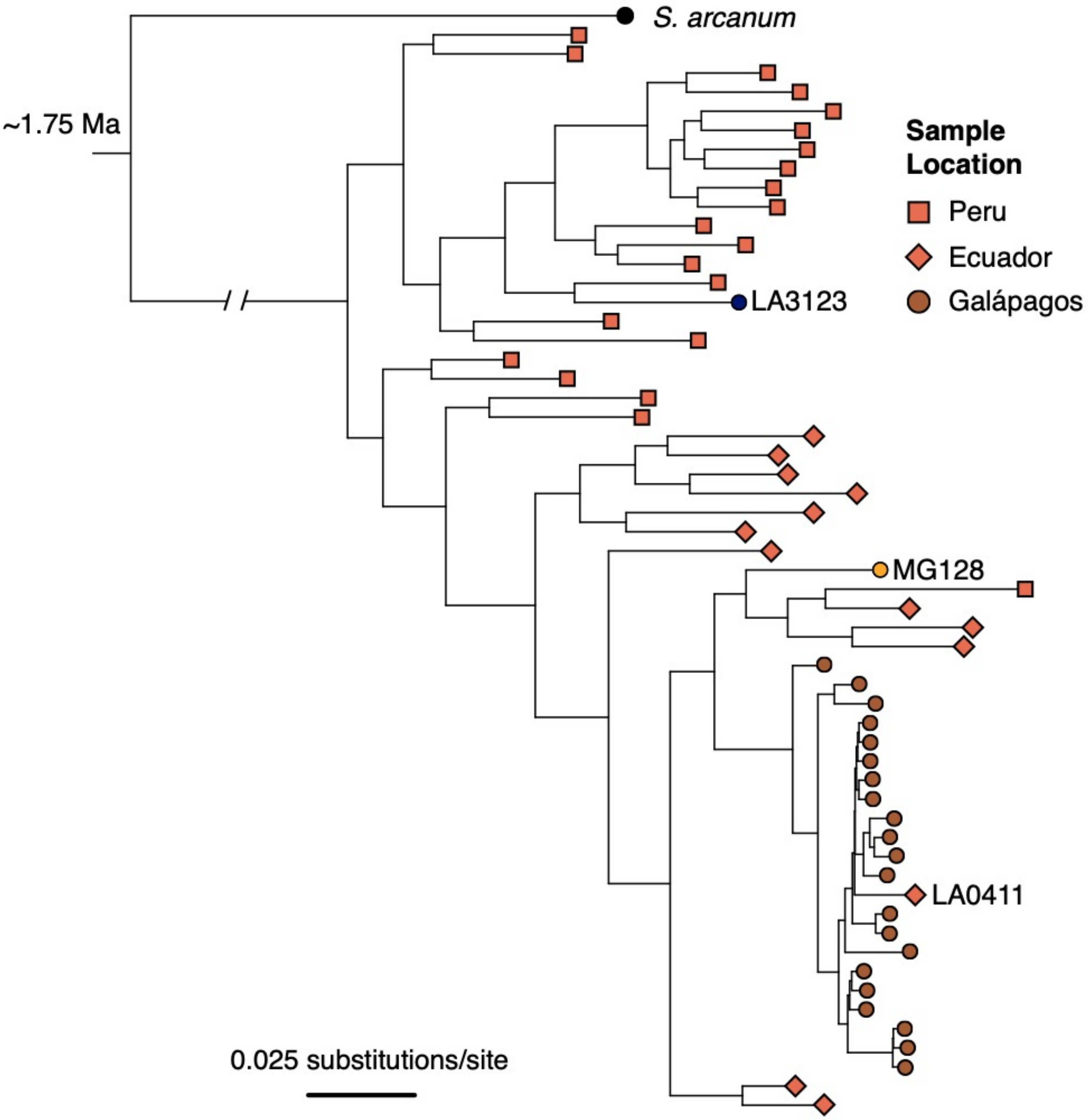


**Figure S3**: ML tree of individual samples inferred with RAxML, using data concatenated across all RAD loci. Samples were subset by population (for Galápagos collections, 1-2 individuals/population) and by geographic region (for mainland accessions; 20 individuals from Peru, 14 individuals from Ecuador) to limit redundancy and increase computation speed.

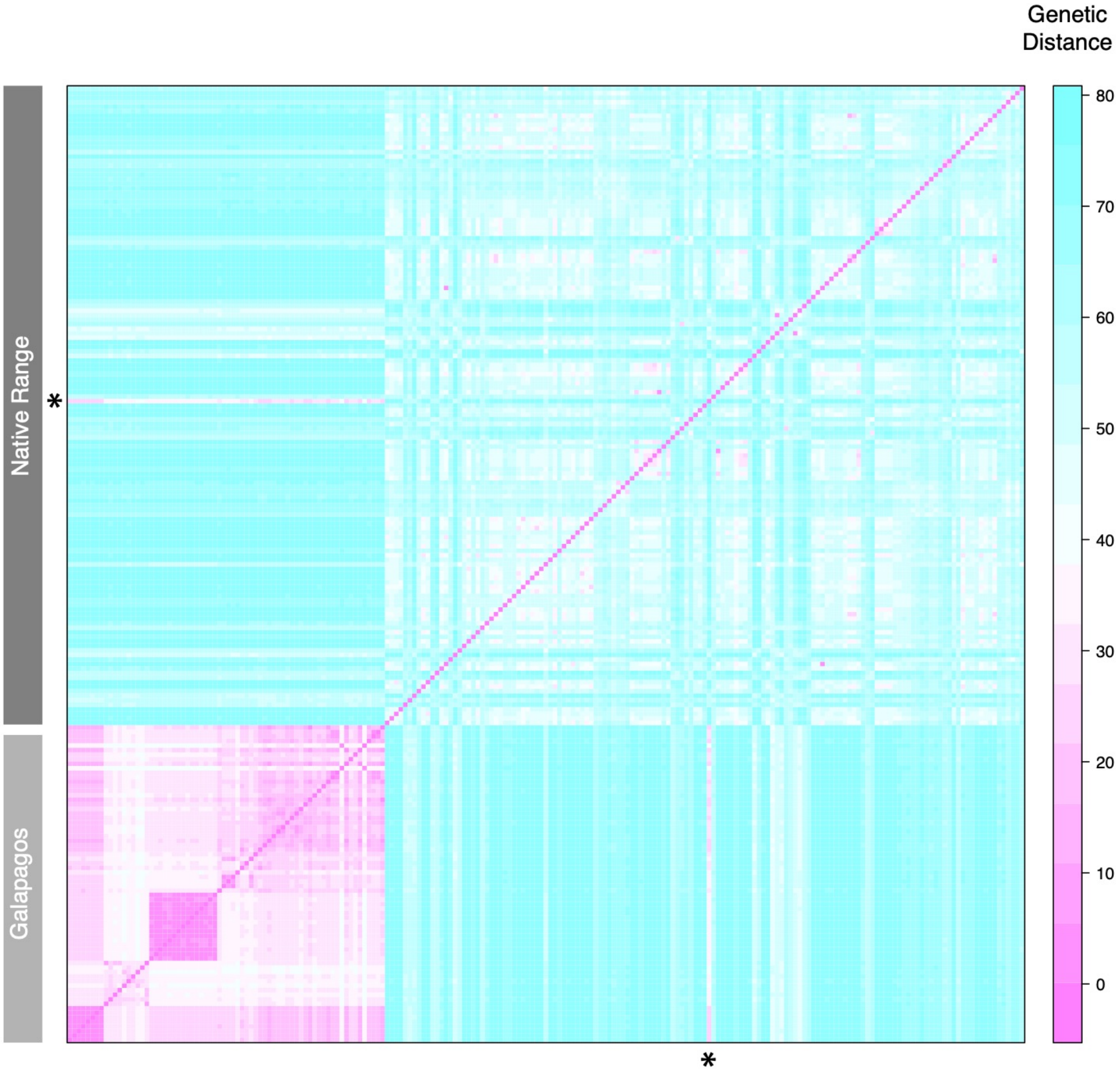


**Figure S4**: Pairwise genetic distances at all polymorphic loci. Row indicated with an asterisk is LA0411, a sample putatively reintroduced to mainland Ecuador from Galápagos.

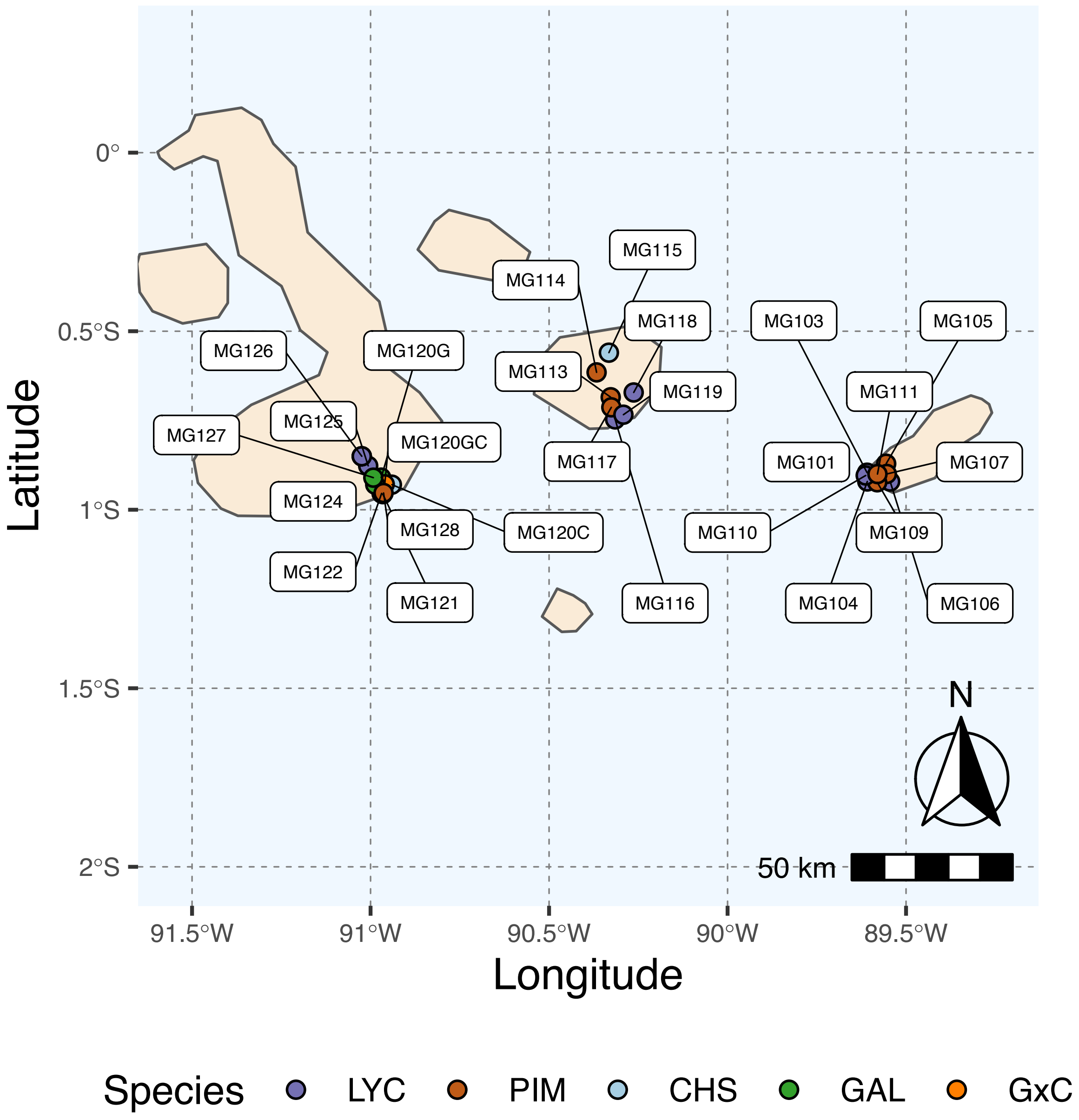


**Figure S5**: Map of all island collection locations. Refer to Table S1 for details.

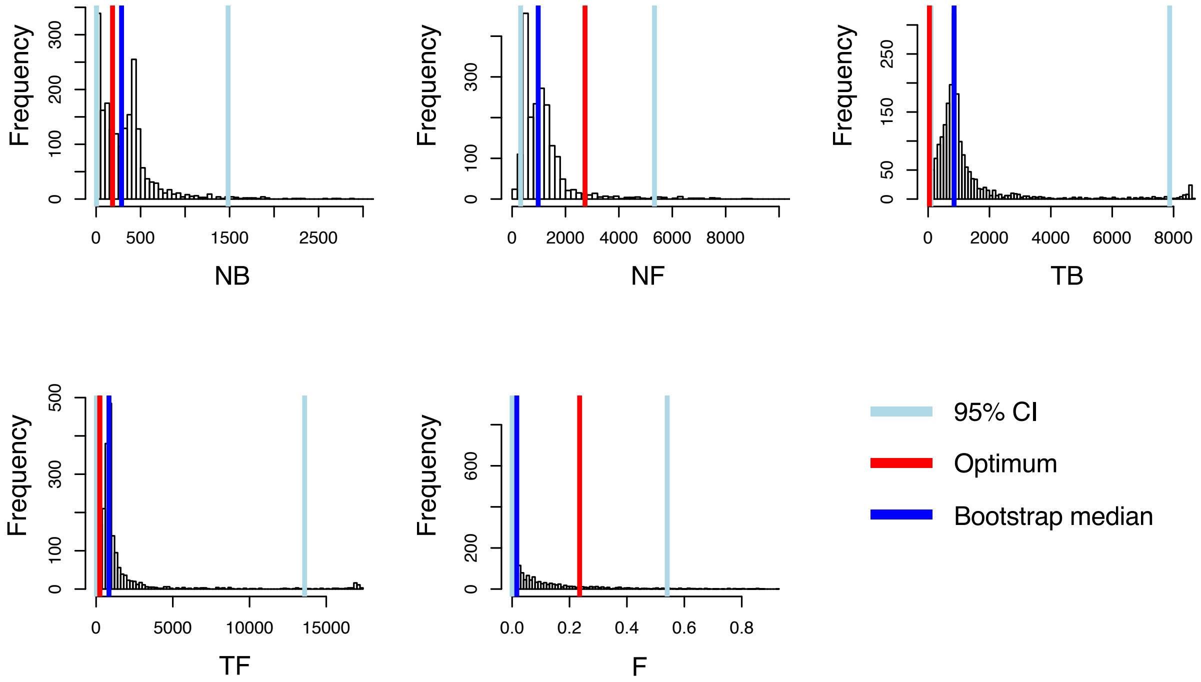


**Figure S6:** Histograms of bootstrapped parameter estimates from *dadi* for PIM population MG114, using the introgression-masked site frequency spectrum. Light blue bars indicate 2.5% and 97.5% quantiles. Red bars indicate the optimum value inferred. Dark blue bars indicate the bootstrapped median value.

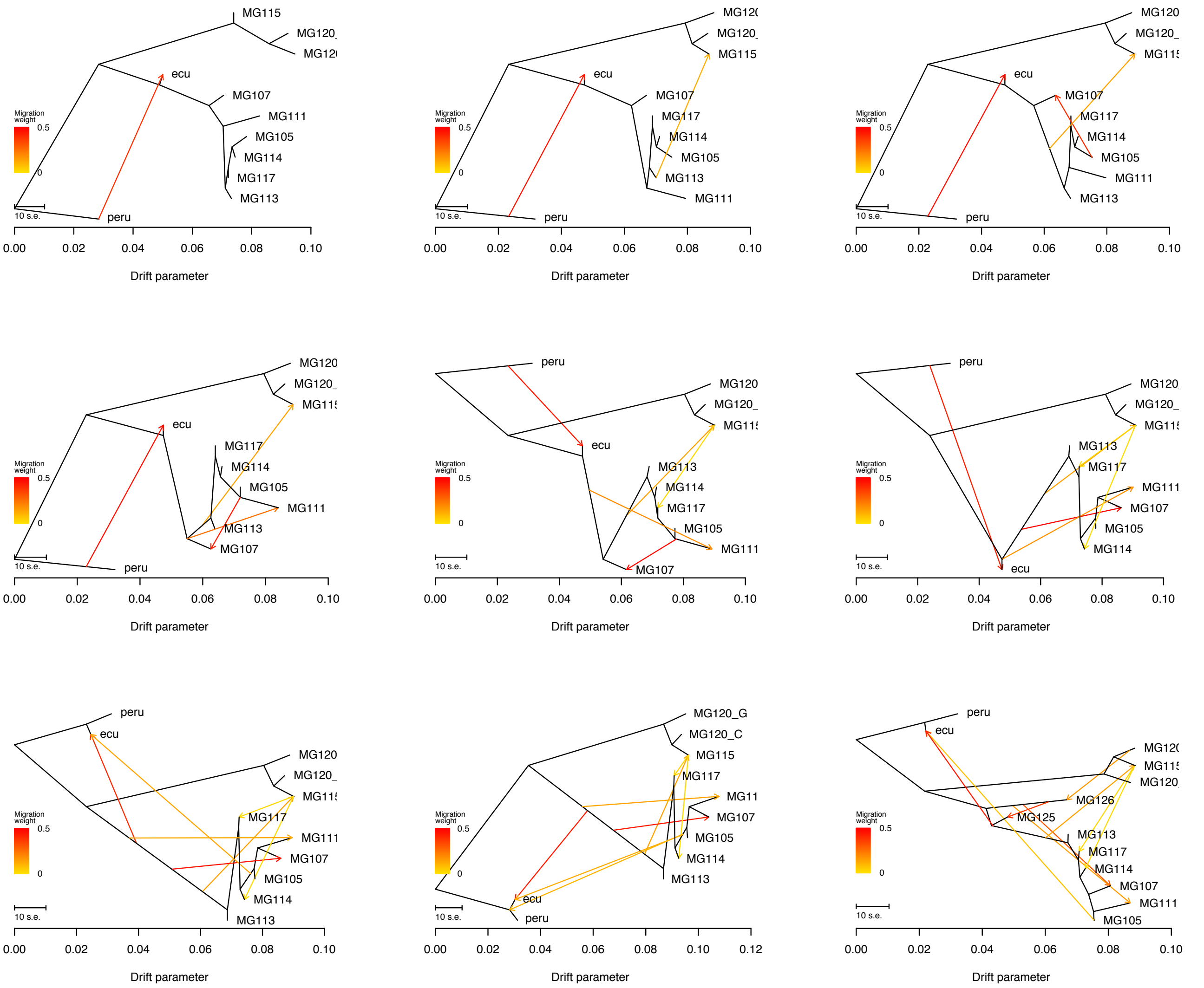


**Figure S7:** *Treemix* summary figures for all tested values for m (migration events; 1-8).

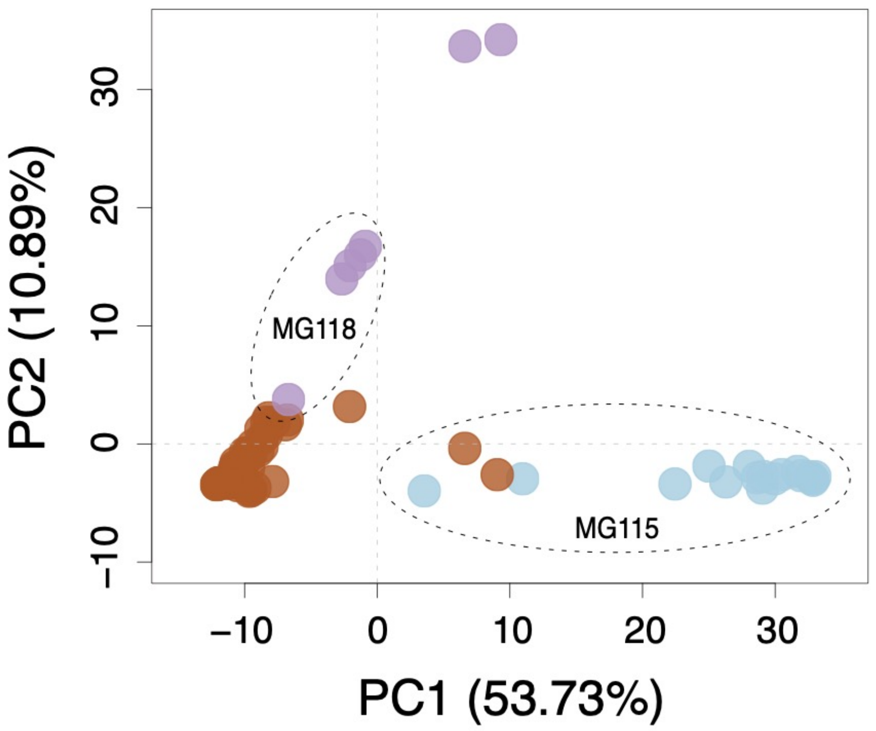


**Figure S8**: Multilocus PCA for Santa Cruz collections. Brown, purple, and blue points correspond to PIM, LYC, and CHS, respectively.

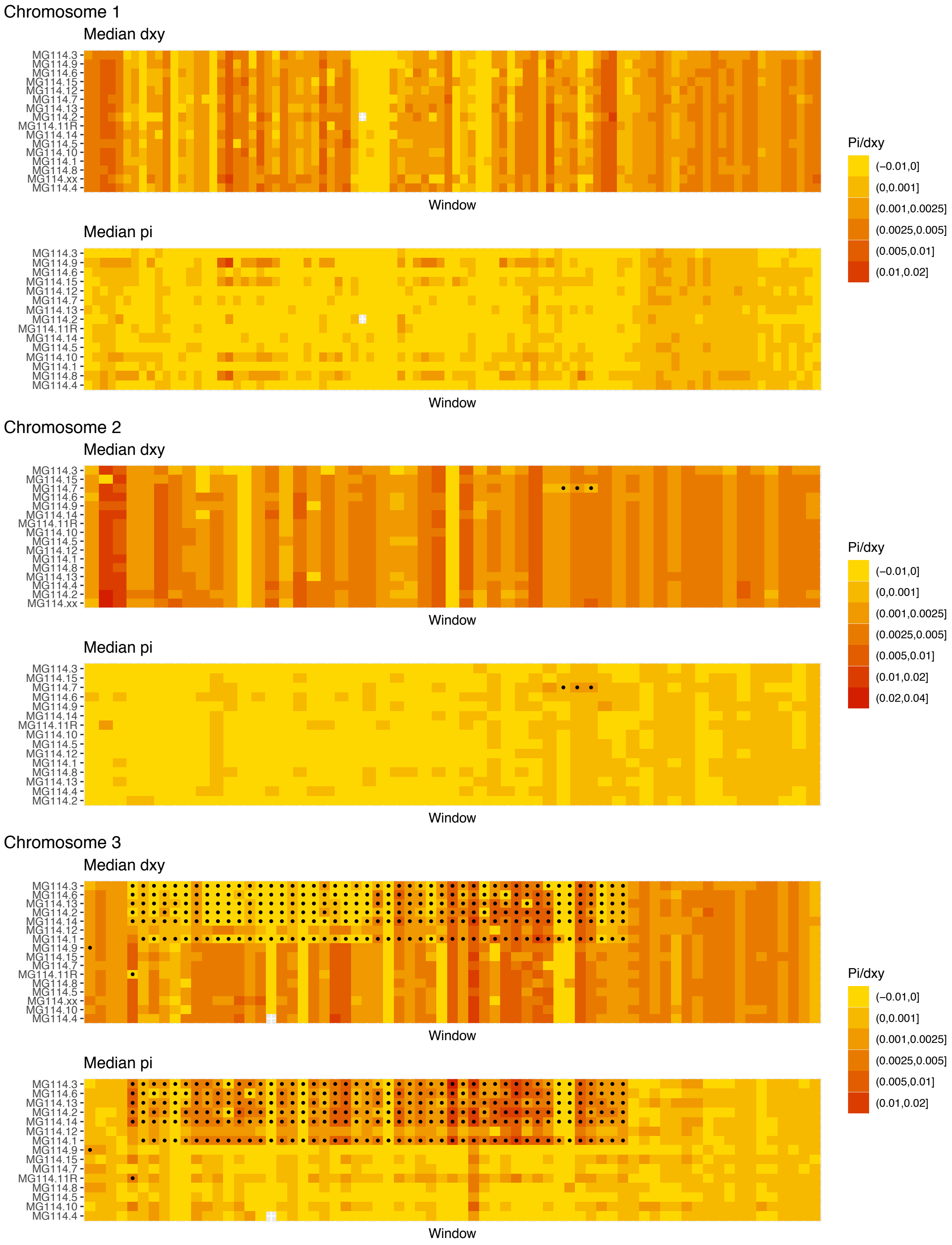
Chromosome 1
Median dxy
MG114.3
MG114.9
MG114.6
MG114.15
MG114.12
MG114.7
MG114.13
MG114.2
MG114.11R
MG114.14
MG114.5
MG114.10
MG114.1
MG114.8
MG114.xx
MG114.4
Window
Median pi
MG114.3
MG114.9
MG114.6
MG114.15
MG114.12
MG114.7
MG114.13
MG114.2
MG114.11R
MG114.14
MG114.5
MG114.10
MG114.1
MG114.8
MG114.4
Window
Pi/dxy
(−0.01,0]
(0,0.001]
(0.001,0.0025]
(0.0025,0.005]
(0.005,0.01]
(0.01,0.02]
Chromosome 2
Median dxy
MG114.3
MG114.15
MG114.7
MG114.6
MG114.9
MG114.14
MG114.11R
MG114.10
MG114.5
MG114.12
MG114.1
MG114.8
MG114.13
MG114.4
MG114.2
MG114.xx
Window
Median pi
MG114.3
MG114.15
MG114.7
MG114.6
MG114.9
MG114.14
MG114.11R
MG114.10
MG114.5
MG114.12
MG114.1
MG114.8
MG114.13
MG114.4
MG114.2
Window
Pi/dxy
(−0.01,0]
(0,0.001]
(0.001,0.0025]
(0.0025,0.005]
(0.005,0.01]
(0.01,0.02]
(0.02,0.04]
Chromosome 3
Median dxy
MG114.3
MG114.6
MG114.13
MG114.2
MG114.14
MG114.12
MG114.1
MG114.9
MG114.15
MG114.7
MG114.11R
MG114.8
MG114.5
MG114.xx
MG114.10
MG114.4
Window
Median pi
MG114.3
MG114.6
MG114.13
MG114.2
MG114.14
MG114.12
MG114.1
MG114.9
MG114.15
MG114.7
MG114.11R
MG114.8
MG114.5
MG114.10
MG114.4
Window
Pi/dxy
(−0.01,0]
(0,0.001]
(0.001,0.0025]
(0.0025,0.005]
(0.005,0.01]
(0.01,0.02]

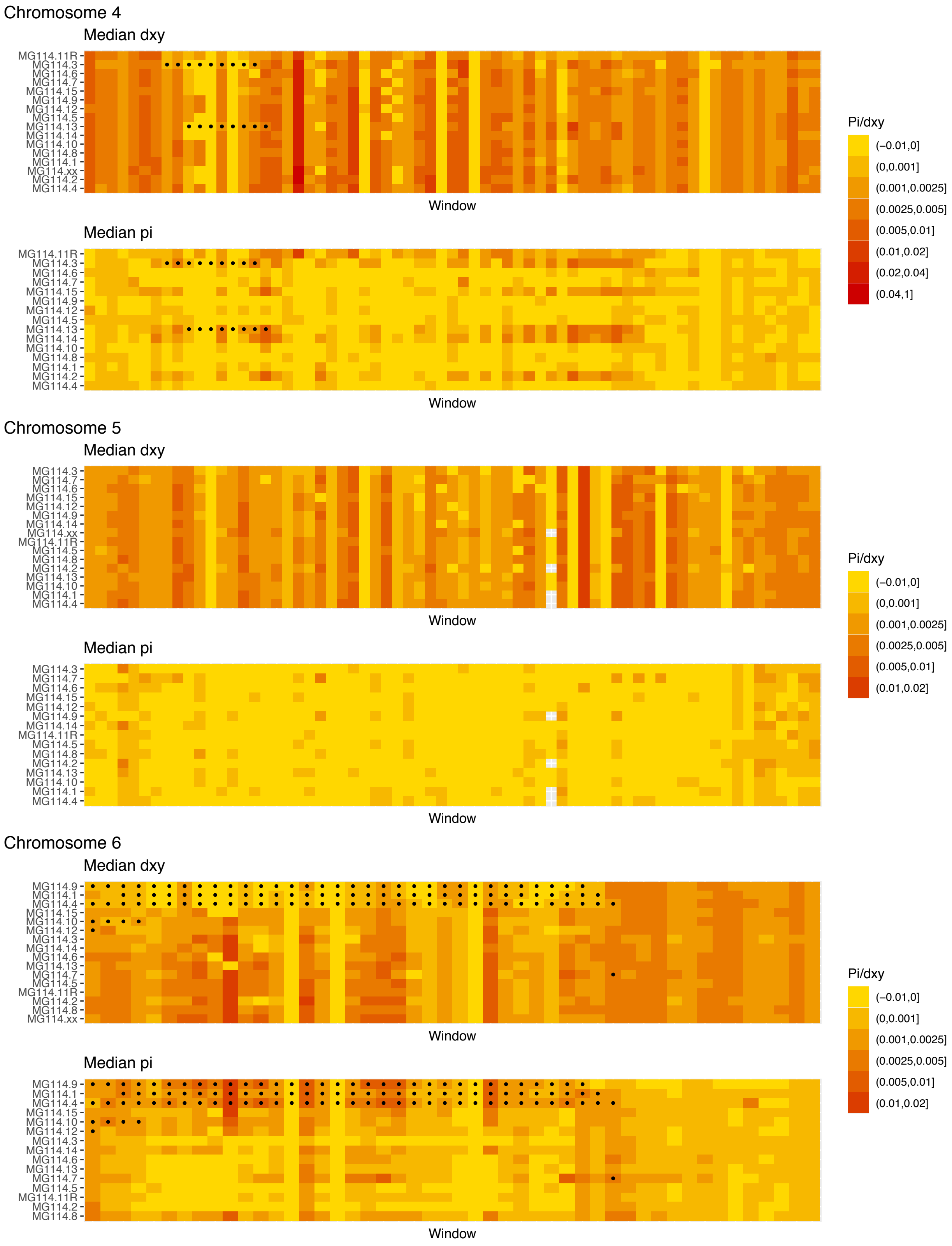
Chromosome 4
Median dxy
MG114.11R
MG114.3
MG114.6
MG114.7
MG114.15
MG114.9
MG114.12
MG114.5
MG114.13
MG114.14
MG114.10
MG114.8
MG114.1
MG114.xx
MG114.2
MG114.4
Window
Median pi
MG114.11R
MG114.3
MG114.6
MG114.7
MG114.15
MG114.9
MG114.12
MG114.5
MG114.13
MG114.14
MG114.10
MG114.8
MG114.1
MG114.2
MG114.4
Window
Pi/dxy
(−0.01,0]
(0,0.001]
(0.001,0.0025]
(0.0025,0.005]
(0.005,0.01]
(0.01,0.02]
(0.02,0.04]
(0.04,1]
Chromosome 5
Median dxy
MG114.3
MG114.7
MG114.6
MG114.15
MG114.12
MG114.9
MG114.14
MG114.xx
MG114.11R
MG114.5
MG114.8
MG114.2
MG114.13
MG114.10
MG114.1
MG114.4
Window
Median pi
MG114.3
MG114.7
MG114.6
MG114.15
MG114.12
MG114.9
MG114.14
MG114.11R
MG114.5
MG114.8
MG114.2
MG114.13
MG114.10
MG114.1
MG114.4
Window
Pi/dxy
(−0.01,0]
(0,0.001]
(0.001,0.0025]
(0.0025,0.005]
(0.005,0.01]
(0.01,0.02]
Chromosome 6
Median dxy
MG114.9
MG114.1
MG114.4
MG114.15
MG114.10
MG114.12
MG114.3
MG114.14
MG114.6
MG114.13
MG114.7
MG114.5
MG114.11R
MG114.2
MG114.8
MG114.xx
Window
Median pi
MG114.9
MG114.1
MG114.4
MG114.15
MG114.10
MG114.12
MG114.3
MG114.14
MG114.6
MG114.13
MG114.7
MG114.5
MG114.11R
MG114.2
MG114.8
Window
Pi/dxy
(−0.01,0]
(0,0.001]
(0.001,0.0025]
(0.0025,0.005]
(0.005,0.01]
(0.01,0.02]

#### Chromosome 7

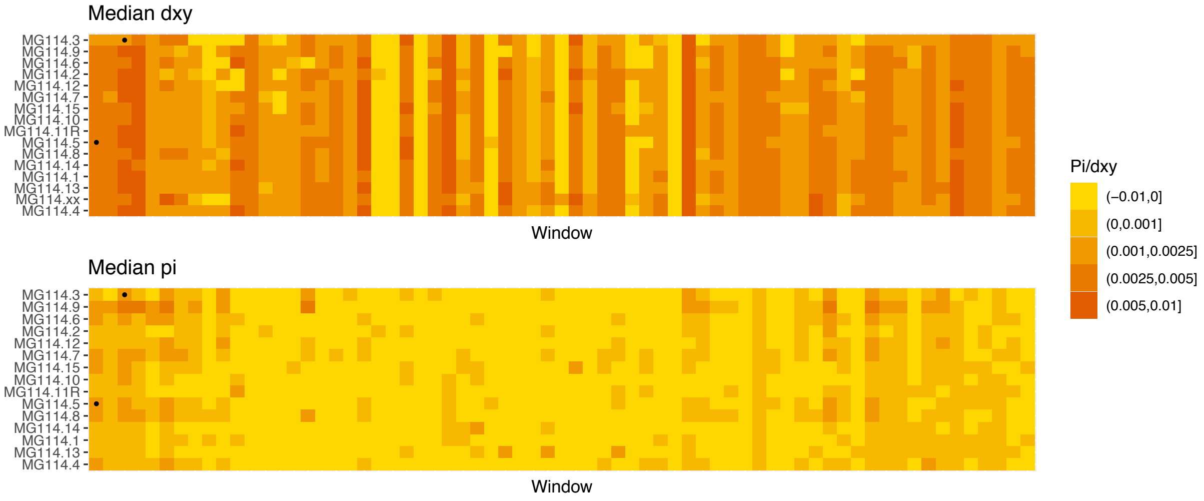


#### Chromosome 8

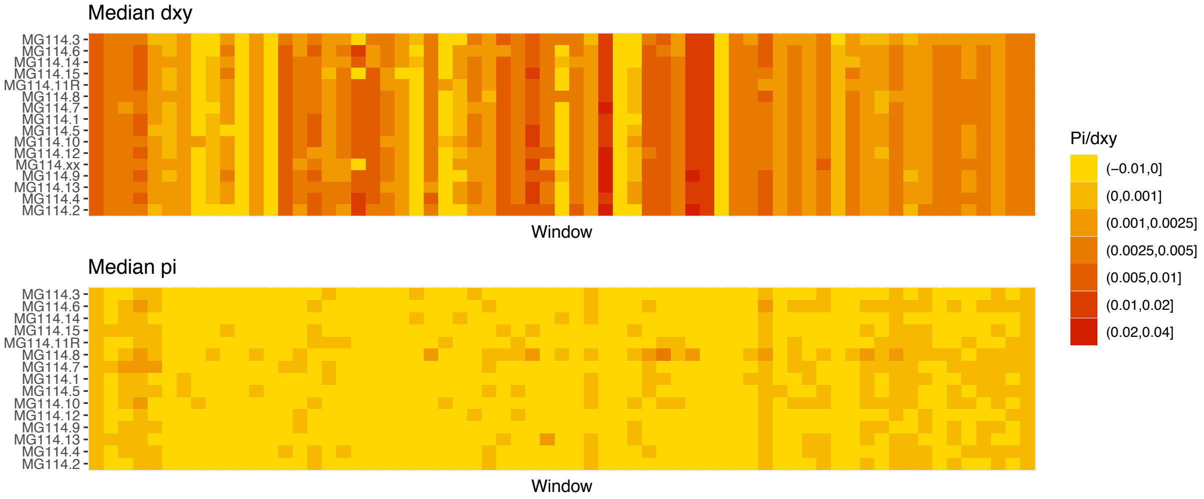


#### Chromosome 9

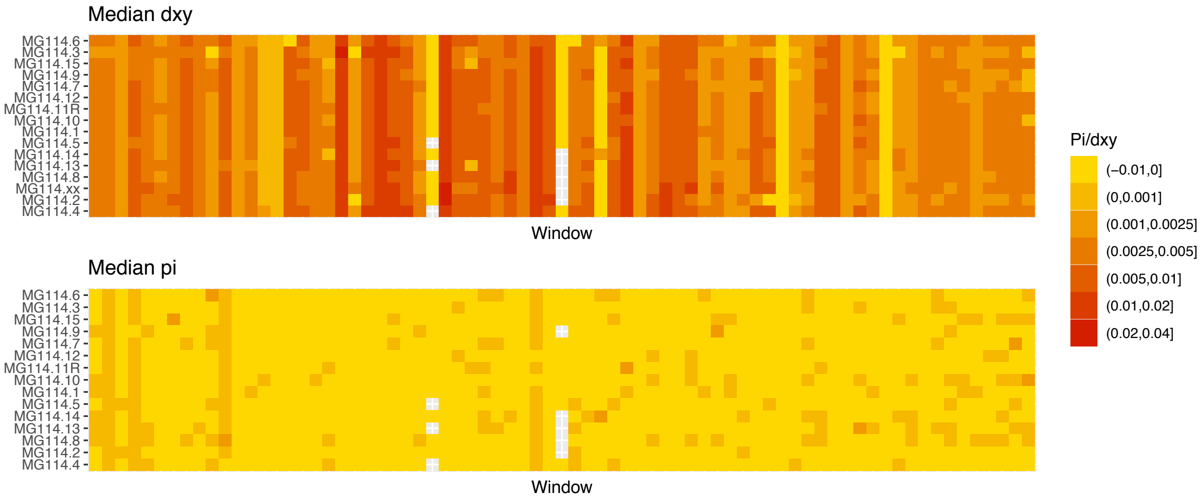

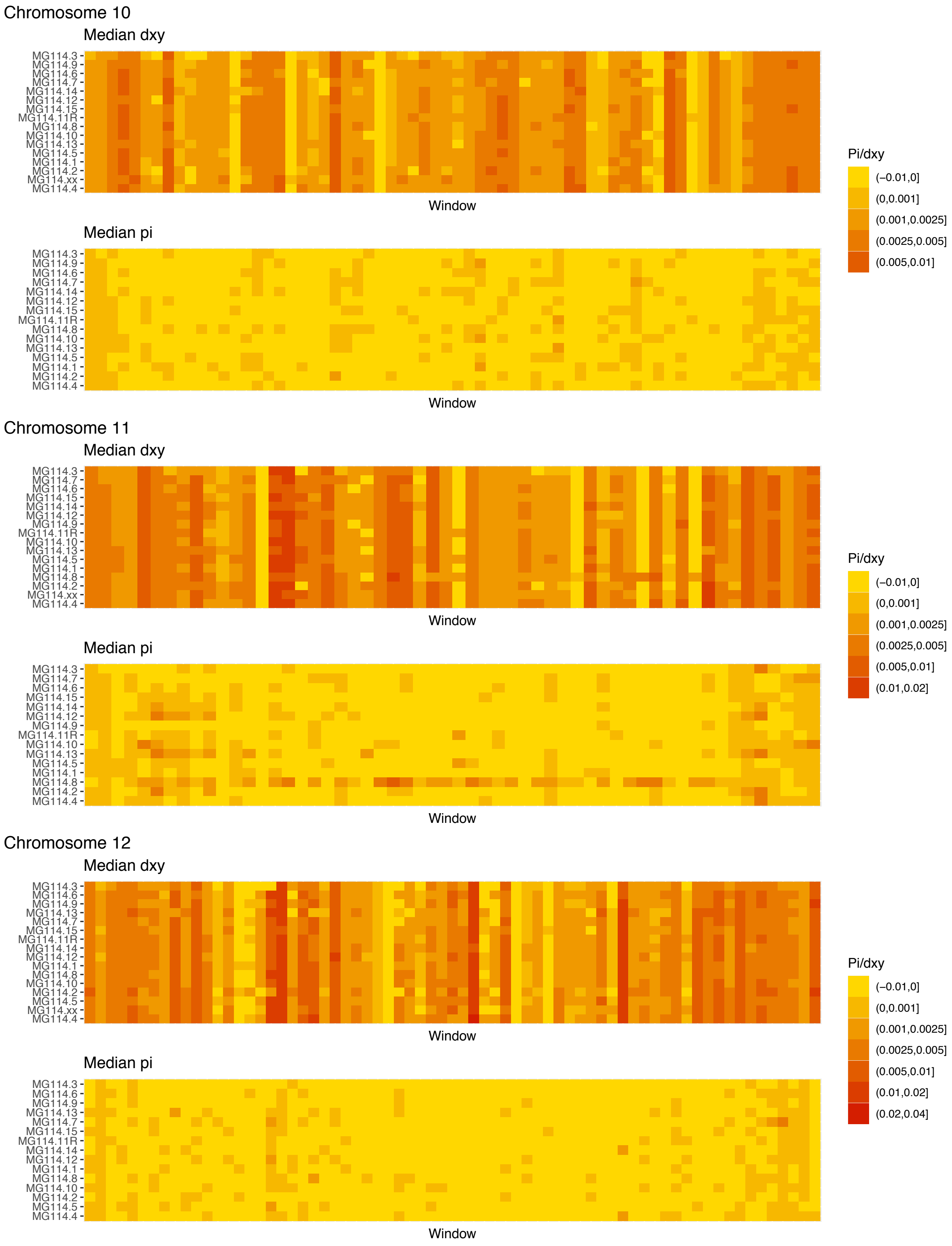


**Figure S9**: Diversity and divergence across the genomes of MG114 individuals. Each cell represents 1 Mb and colors correspond to median π or $d_{XY}$ (to CHS population MG115). Black dots show regions of CHS ancestry predicted by the HMM. HMM predictions were done in 100kb windows. Annotations here reflect the consensus prediction of all windows within each 1 Mb cell.

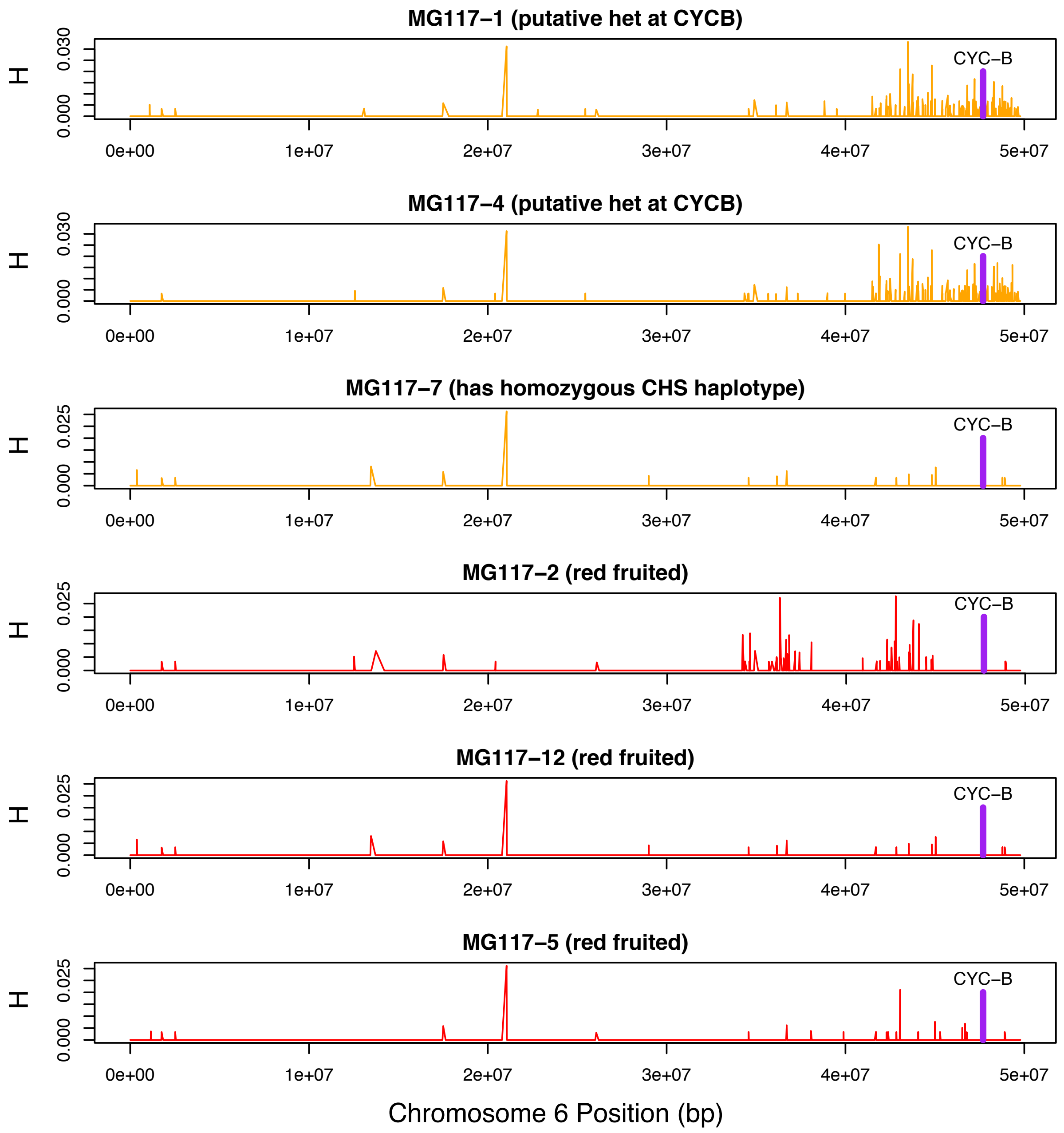


**Figure S10**: Heterozygosity along chromosome 6 of population MG117. CHS ancestry at *CYC-B* (dashed line) in MG117-1 and MG117-4 is heterozygous, as shown by elevated heterozygosity estimates at that location in these individuals.

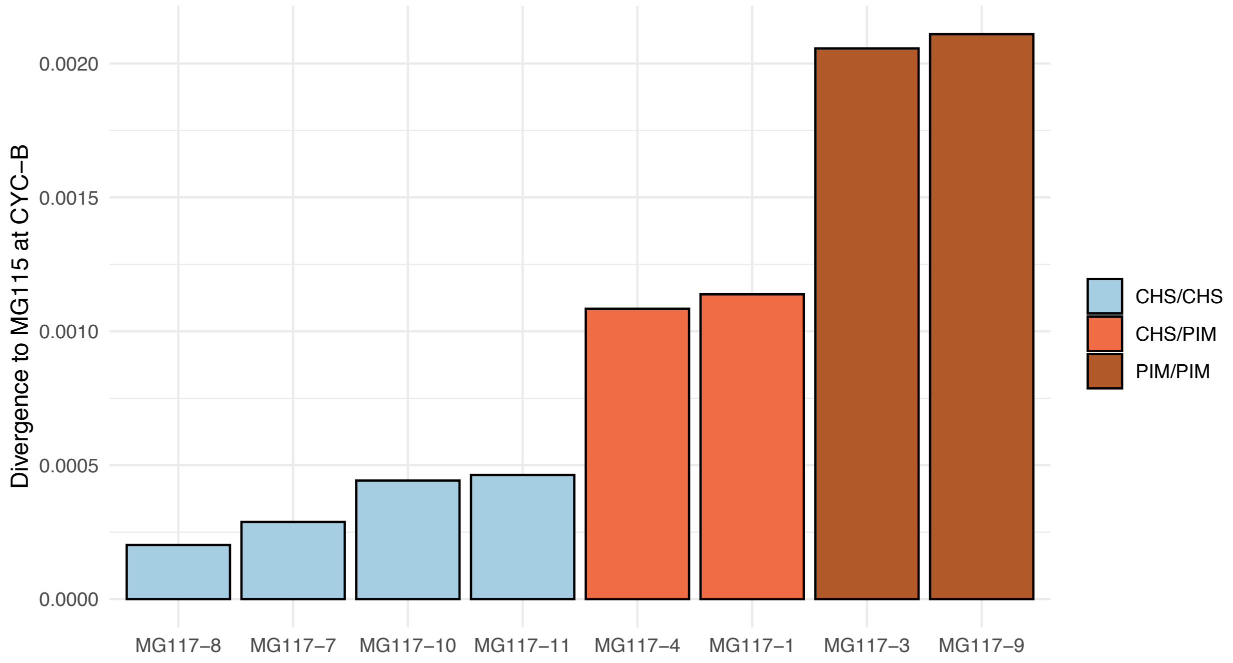


**Figure S11:** Median divergence estimates between MG117 individuals and MG115 at CYC-B. The HMM correctly classifies intermediately diverged regions as heterozygous.

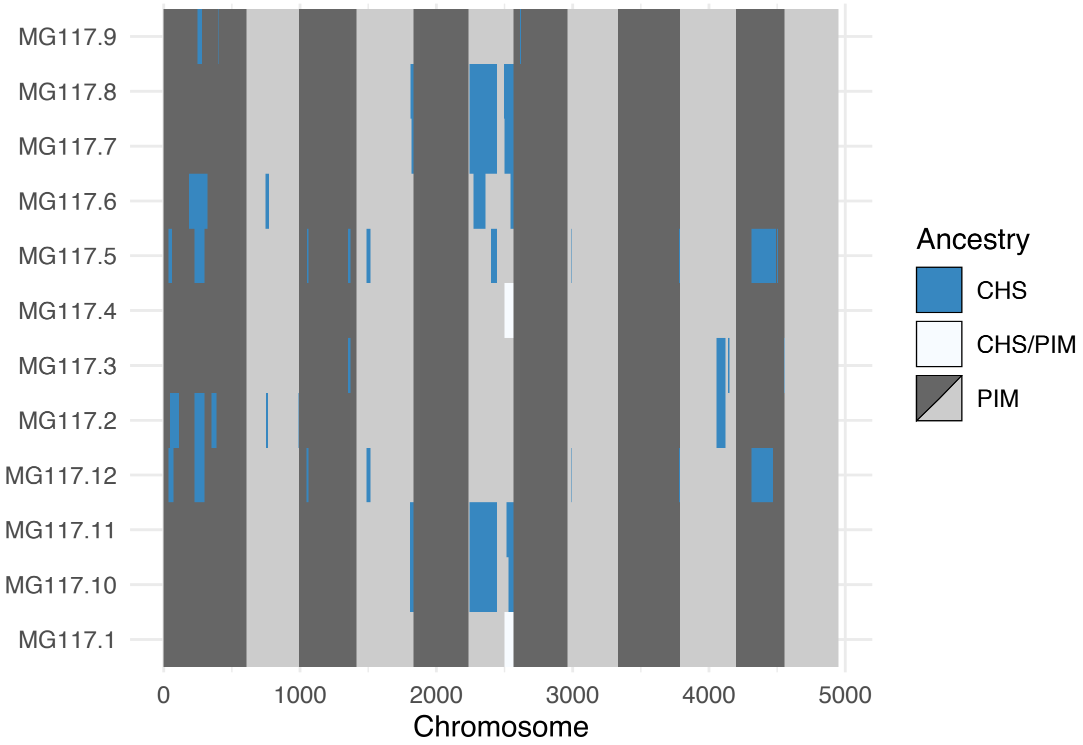


**Figure S12**: Genome-wide local ancestry in population MG117 as inferred by the HMM.

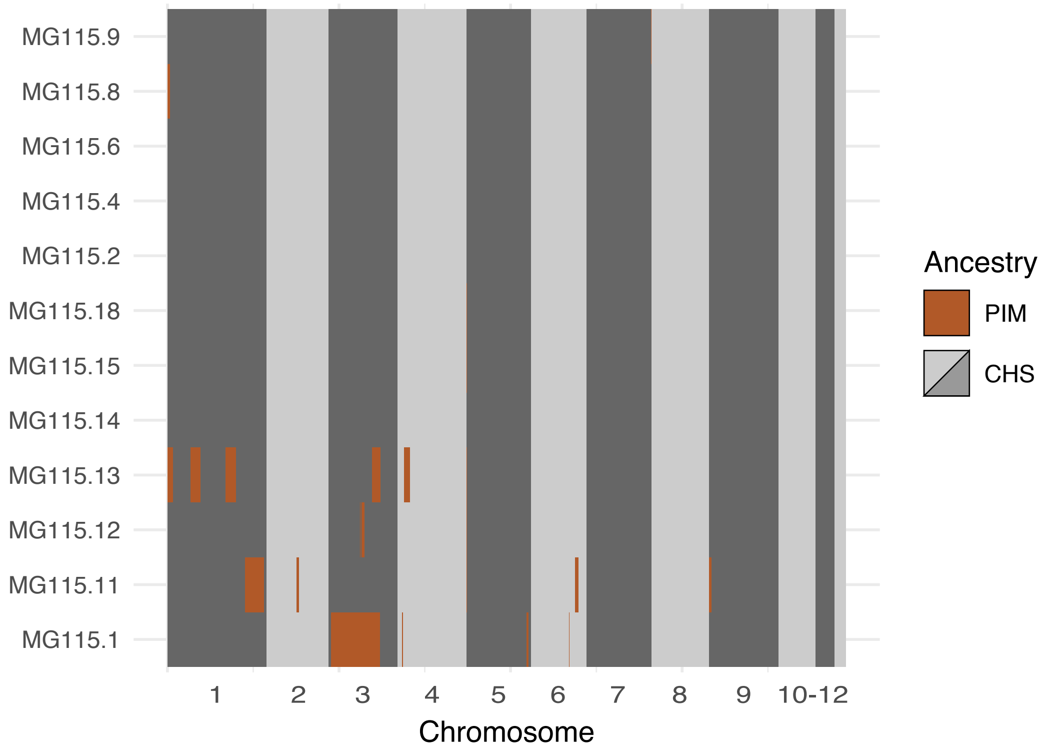


**Figure S13**: Local ancestry assignment throughout the genomes of CHS plants from population MG115 (Santa Cruz), using MG114 as the PIM reference population. These data represent the opposite direction of gene flow (PIM → CHS) from that shown in Figure 5B (CHS → PIM). Admixture in the PIM → CHS direction was markedly lower than CHS → PIM.

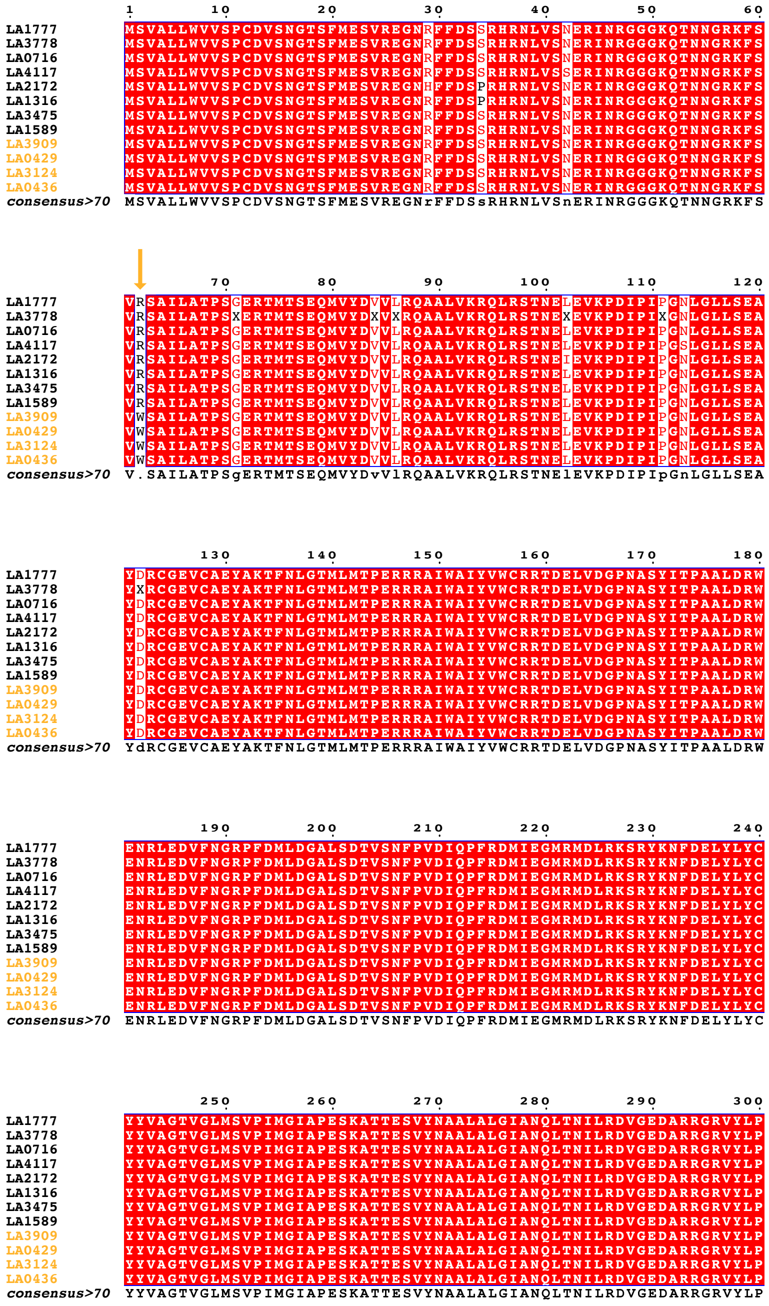

**Figure S14**: Coding sequence alignment of *PSY1* for 9 wild tomato species (12 accessions). Endemic accessions are colored orange. The nonsynonymous substitution which defines the endemic clade is indicated with an orange arrow. Data were obtained from Pease et al. (2016).

**Table S1**: Full list of population collection sites and their geographic coordinates.

| Island | Population ID | Species | Elevation (m) | Lat | Lon | Description |
|---|---|---|---|---|---|---|
| San Cristobal | MG105 | PIM | 319 | 00° 54' 28.76" S | 89° 33' 20.37" W | El Progresso |
| | MG107 | PIM | 319 | 00° 53' 58.99" S | 89° 33' 13.66" W | Soledad |
| | MG111 | PIM | 8 | 00° 54' 04.09" S | 89° 36' 36.28" W | Puerto Baquerizo |
| Santa Cruz | MG113 | PIM | 259 | 00° 41' 05.56" S | 90° 19' 38.02" W | Bellavista |
| | MG114 | PIM | 571 | 00° 36' 54.89" S | 90° 22' 00.64" W | Mina Roja |
| | MG116 | PIM/LYC | 10 | 00° 44' 49.02" S | 90° 18' 53.42" W | Puerto Ayora |
| | MG117 | PIM | 129 | 00° 42' 47.28" S | 90° 19' 31.81" W | Thomas Berlanga |
| | MG118 | PIMxLYC | 264 | 00° 40' 17.50" S | 90° 15' 46.42" W | Cascajo |
| | MG115 | CHS | 251 | 00° 33' 36.23" S | 90° 19' 56.65" W | Santa Cruz Hwy |
| Isabela | MG120G | GAL | 9 | 00° 55' 45.81" S | 90° 58' 44.25" W | Laguna Manzanilla |
| | MG120C | CHS | 9 | 00° 55' 45.81" S | 90° 58' 44.25" W | Laguna Manzanilla |
| | MG124 | GAL | 12 | 00° 55' 47.27" S | 90° 59' 12.43" W | Road to Santa Tomás |
| | MG127 | GAL | 16 | 00° 55' 23.01" S | 90° 59' 20.93" W | Road to Santa Tomás |
| | MG122 | LYC | 6 | 00° 57' 15.21" S | 90° 58' 08.41" W | Puerto Villamil |
| | MG125 | LYC | 138 | 00° 52' 37.26" S | 91° 00' 23.73" W | Puerto Villamil |
| | MG126 | LYC | 332 | 00° 51' 00.49" S | 91° 01' 28.39" W | Santa Tomás |
| | MG128 | PIM | 7 | 00° 57' 11.17" S | 90° 57' 48.62" W | Puerto Villamil |

**Table S2**: List of mainland collection sites

| Accession | Longitude | Latitude |
|---|---|---|
| LA0121 | -78.99556 | -8.18444 |
| LA0122 | -78.72 | -8.01 |
| LA0373 | -78.218333 | -9.936944 |
| LA0375 | -78.21 | -9.94 |
| LA0418 | -79.93333 | -1.86667 |
| LA 0420 | -79.65 | -1.06 |
| LA 0442 | -78.25917 | -9.48167 |
| LA 0443 | -79.48333 | -1.1 |
| LA 0753 | -79.03333 | -8.11667 |
| LA 0722 | -76.88333 | -12.26667 |
| LA 1248 | -79.35 | -3.98333 |
| LA 1256 | -79.61667 | -2.66667 |
| LA 1280 | -76.71667 | -12.03333 |
| LA 1301 | -75.91667 | -13.73333 |
| LA 1381 | -79.96667 | -5.56667 |
| LA 1416 | -79.333333 | -0.266667 |
| LA 1428 | -80.216667 | -0.816667 |
| LA 1469 | -79.79 | -5.86 |
| LA 1470 | -79.73333 | -6.06667 |
| LA 1472 | -76.25 | -13.31667 |
| LA 1478 | -80.08333 | -5.21667 |
| LA 1520 | -77.11889 | -11.04528 |
| LA 1547 | -77.93333 | 0.58333 |
| LA 1561 | -77.41667 | -11.1 |
| LA 1562 | -76.81667 | -12.13333 |
| LA 1572 | -76.78333 | -11.96667 |
| LA 1576 | -76.86667 | -12.16667 |
| LA 1579 | -79.87 | -6.59 |
| LA 1584 | -79.79 | -6.37 |
| LA 1586 | -78.73 | -8.36 |
| LA 1589 | -78.74 | -8.39 |
| LA 1598 | -78.19 | -9.93 |
| LA 1600 | -78.04 | -10.28 |
| LA 1601 | -77.68 | -10.67 |
| LA 1602 | -76.98333 | -11.78333 |
| LA 1603 | -77 | -11.53333 |
| LA 1604 | -77.08333 | -11.48333 |
| LA 1605 | -76.41667 | -13.08333 |
| LA 1611 | -76.61667 | -12.6 |
| LA 1614 | -77.36667 | -11.11667 |
| LA 1615 | -80.63333 | -5.23333 |
| LA 1635 | -75.06667 | -14.63333 |
| LA 1636 | -76.08333 | -13.45 |
| LA 1637 | -75.83333 | -13.36667 |
| LA 1651 | -76.94611 | -12.08528 |
| LA 1659 | -77.85861 | -9.54667 |
| LA 1660 | -77.96667 | -9.53333 |
| LA 1661 | -76.5 | -12.76667 |
| LA 1670 | -70.51667 | -17.83333 |
| LA 1683 | -81.11 | -4.87 |
| LA 1684 | -80.15 | -5.1 |
| LA 1685 | -80.6975 | -4.88667 |
| LA 1686 | -80.62 | -5.07 |
| LA 1687 | -80.62 | -5.07 |
| LA 1688 | -80.375 | -4.88333 |
| LA 1689 | -80.6175 | -5.17639 |
| LA 1690 | -80.6175 | -5.17639 |
| LA 1697 | -76.98333 | -12.05 |
| LA 1728 | -75.95 | -13.4 |
| LA 1729 | -75.64056 | -13.29694 |
| LA 1921 | -75.12722 | -14.31194 |
| LA 1923 | -75.28333 | -14.66667 |

| | | |
|---|---|---|
| LA 1924 | -75.21417 | -14.62889 |
| LA 1933 | -74.44583 | -15.45639 |
| LA 1936 | -74.0325 | -15.83361 |
| LA 1950 | -73.25806 | -16.38861 |
| LA 1987 | -78.83 | -8.43 |
| LA 2069 | -79.649722 | -4.050556 |
| LA 2093 | -79.88806 | -3.62806 |
| LA 2102 | -79.4675 | -4.40167 |
| LA 2181 | -78.78306 | -5.77583 |
| LA 2186 | -78.16667 | -5.89167 |
| LA 2401 | -78.22778 | -9.50833 |
| LA 2412 | -76.71667 | -12.48333 |
| PI 251318 | -79.964167 | -5.260833 |
| PI 251319 | -79.48333 | -1.1 |
| PI 251320 | -79.48333 | -1.1 |
| PI 251321 | -80.26667 | -2.33333 |
| LA 2544 | -79.33333 | -7.3 |
| LA 2576 | -77.89167 | -9.55 |
| LA 2578 | -77.925 | -9.525 |
| LA 2649 | -80.16667 | -5.575 |
| LA 2725 | -75.86944 | -13.71833 |
| LA 2832 | -75.13333 | -14.46667 |
| LA 2839 | -78.06667 | -5.925 |
| LA 2852 | -80.48333 | -0.83333 |
| LA 2854 | -80.58333 | -1.35 |
| LA 2857 | -90.96667 | -0.95 |
| LA 2915 | -79.74528 | -5.98472 |
| LA 2933 | -80.5625 | -1.44167 |
| LA 2983 | -76.76278 | -12.53194 |
| LA 3123 | -90.38306 | -0.62333 |
| PI 365909 | -79.15 | -0.25 |
| PI 365910 | -79.85 | 0.86667 |
| PI 365911 | -79.96667 | -3.45833 |
| PI 365912 | -79.36 | -3.99 |
| PI 365914 | -79.46667 | -1.58333 |
| PI 365915 | -79.53333 | -1.55 |
| PI 365916 | -79.51667 | -1.81667 |
| PI 365917 | -79.63 | -2.25 |
| PI 365918 | -79.816667 | -2.066667 |
| PI 365958 | -74.95 | -14.83333 |
| PI 365960 | -78.32944 | -9.46028 |
| PI 365961 | -78.68056 | -8.53639 |
| PI 365962 | -78.88806 | -8.08611 |
| PI 365964 | -78.35 | -9.18333 |
| PI 365965 | -78.15 | -9.04167 |
| PI 365966 | -77.9 | -9.55833 |
| PI 379021 | -78.66667 | -7.98333 |
| PI 379023 | -76.65 | -11.85833 |
| PI 379024 | -76.65389 | -11.88944 |
| PI 379025 | -75.15833 | -14.68333 |
| PI 379026 | -80.05056 | -5.2525 |
| PI 379059 | -80.1 | -1.48333 |
| PI 390519 | -79.9833 | -3.55 |
| PI 390688 | -77.633 | -10.333 |
| PI 390689 | -77.733 | -10.5 |
| PI 390690 | -77.5 | -1.1 |
| PI 390691 | -77.766 | -10.633 |
| PI 390695 | -79.6166 | -7 |
| PI 390696 | -79.6166 | -7 |
| PI 390699 | -79.9 | -6.68 |
| PI 390700 | -79.7 | -6.1666 |
| PI 390701 | -79.91666 | -5.6 |
| PI 390705 | -80.25 | -5 |
| PI 390706 | -80.75 | -5.5 |
| PI 390707 | -80.816667 | -5.5333 |
| PI 390716 | -81.066666 | -4.8833333 |
| PI 390718 | -79.96666 | -5.266666 |
| PI 390721 | -79.71666 | -5.416667 |
| PI 390722 | -80.75 | -4.85 |
| PI 390728 | -80.51666 | -3.616666 |
| PI 390748 | -80.416666 | -3.55 |
| PI 407533 | -76.71 | -11.95 |
| PI 407534 | -76.83333 | -11.98333 |
| PI 407535 | -76.76667 | -12.08333 |
| PI 407537 | -79.87 | -6.59 |

| | | |
|---|---|---|
| PI 407538 | -79.89 | -6.6 |
| PI 407544 | -79 | -8.10833 |
| PI 407550 | -78.26 | -9.49 |
| PI 407551 | -78.47 | -9.27 |
| PI 407552 | -78.56667 | -8.925 |
| PI 407553 | -78.34 | -9.47 |
| PI 407554 | -78.126389 | -10.067222 |
| PI 503517 | -80.18333 | -5.16667 |
| PI 503519 | -79.98333 | -5.175 |
| PI 503521 | -80.58333 | -4.75 |
| PI 503522 | -80.825 | -4.90833 |
| PI 503523 | -80.625 | -5.21667 |
| PI 503524 | -80.7 | -3.8 |

**Table S3**: Stacks ref_map assembly summary

| Statistic | Value |
|---|---|
| Bam records | 444,288,642 |
| Primary alignments retained | 688,775,808 |
| Records per sample | 2,128,095 (17,586-13,836,425) |
| Assembled/genotyped loci | 269,892 |
| Effective per-sample coverage | 61.4x (sd = 35.0x) |
| Mean sites per locus | 200.3 |

**Table S4**: Stacks per-sample assembly summary

| Sample | Bam records | Fraction kept | # Loci | Mean coverage |
|---|---|---|---|---|
| 0121-101 | 860527 | 0.974 | 17034 | 33 |
| 0122-101 | 1630820 | 0.972 | 25079 | 48.589 |
| 0373-102 | 3329913 | 0.972 | 23171 | 103.581 |
| 0375-101 | 2126539 | 0.966 | 32746 | 54.023 |
| 0418-101 | 2143602 | 0.978 | 32138 | 55.606 |
| 0420-101 | 1719200 | 0.979 | 29998 | 46.226 |
| 0442-101 | 4607456 | 0.975 | 46057 | 102.695 |
| 0443-101 | 2497478 | 0.973 | 35378 | 61.342 |
| 0753-101 | 1107015 | 0.97 | 24562 | 32.976 |
| 0772-101 | 2065633 | 0.972 | 27580 | 57.748 |
| 1248-101 | 2745125 | 0.981 | 30936 | 73.403 |
| 1256-101 | 3632845 | 0.978 | 35536 | 90.728 |
| 1280-101 | 3120897 | 0.977 | 30504 | 83.85 |
| 1301-101 | 2155029 | 0.978 | 31797 | 56.615 |
| 1316-1 | 2454676 | 0.938 | 24096 | 81.476 |
| 1316-6 | 2622282 | 0.939 | 26819 | 82.11 |
| 1381-101 | 2114032 | 0.974 | 33684 | 53.253 |
| 1416-101 | 313218 | 0.98 | 13820 | 13.808 |
| 1428-101 | 2700077 | 0.982 | 35638 | 68.555 |
| 1469-101 | 1457202 | 0.973 | 29690 | 38.898 |
| 1470-101 | 3475525 | 0.975 | 35756 | 86.178 |
| 1472-101 | 12103090 | 0.971 | 45961 | 291.096 |
| 1478-101 | 989959 | 0.978 | 19611 | 34.414 |
| 1520-101 | 4548266 | 0.976 | 29953 | 125.193 |
| 1547-101 | 874210 | 0.981 | 23120 | 27.471 |
| 1561-101 | 2172059 | 0.978 | 27342 | 61.321 |
| 1562-101 | 2288002 | 0.976 | 27199 | 64.904 |
| 1572-101 | 4330309 | 0.978 | 35422 | 110.974 |
| 1576-101 | 4227007 | 0.968 | 38571 | 106.326 |
| 1579-101 | 2739722 | 0.974 | 30155 | 73.004 |
| 1584-101 | 3938382 | 0.973 | 37777 | 95.768 |
| 1586-101 | 3544559 | 0.972 | 37577 | 85.758 |
| 1589-12 | 823915 | 0.972 | 17903 | 30.026 |
| 1598-101 | 4697846 | 0.977 | 33154 | 120.776 |
| 1600-101 | 2096174 | 0.972 | 30537 | 56.59 |
| 1601-101 | 301039 | 0.963 | 11510 | 15.188 |

| | | | | |
|---|---|---|---|---|
| 1602-101 | 2736356 | 0.972 | 35178 | 67.97 |
| 1603-101 | 5257707 | 0.977 | 37743 | 130.705 |
| 1604-101 | 2492691 | 0.946 | 28257 | 67.72 |
| 1605-101 | 3183592 | 0.98 | 37344 | 76.445 |
| 1611-101 | 2652137 | 0.972 | 30520 | 72.3 |
| 1614-101 | 2035135 | 0.978 | 26826 | 57.963 |
| 1615-101 | 2245827 | 0.979 | 28566 | 63.405 |
| 1635-101 | 5004334 | 0.968 | 41560 | 119.504 |
| 1636-101 | 1900561 | 0.976 | 30965 | 49.755 |
| 1637-101 | 1664781 | 0.978 | 25995 | 49.517 |
| 1651-101 | 266205 | 0.98 | 14669 | 11.852 |
| 1659-101 | 5447399 | 0.979 | 41153 | 129.37 |
| 1660-101 | 1923801 | 0.968 | 26594 | 56.287 |
| 1661-101 | 2289988 | 0.979 | 31529 | 60.589 |
| 1670-101 | 3133778 | 0.978 | 31472 | 83.181 |
| 1683-101 | 3318632 | 0.969 | 35756 | 81.26 |
| 1684-101 | 3191539 | 0.975 | 22414 | 105.04 |
| 1685-101 | 2731893 | 0.977 | 31859 | 71.467 |
| 1686-101 | 3440511 | 0.977 | 35442 | 85.595 |
| 1687-101 | 4310773 | 0.979 | 38098 | 103.703 |
| 1688-101 | 4800347 | 0.978 | 40430 | 114.266 |
| 1689-101 | 1127857 | 0.98 | 21617 | 36.639 |
| 1690-101 | 2555961 | 0.976 | 31373 | 69.644 |
| 1697-101 | 5430669 | 0.978 | 41428 | 128.326 |
| 1728-101 | 5326058 | 0.961 | 21262 | 186.408 |
| 1729-101 | 5544196 | 0.98 | 43872 | 129.781 |
| 1777-27 | 1692290 | 0.93 | 19922 | 62.233 |
| 1921-101 | 2499661 | 0.971 | 31127 | 67.084 |
| 1923-101 | 3376990 | 0.974 | 33440 | 86.66 |
| 1924-101 | 3200102 | 0.977 | 33993 | 82.095 |
| 1933-101 | 2357305 | 0.978 | 32867 | 61.884 |
| 1936-101 | 13836425 | 0.975 | 41433 | 332.854 |
| 1950-101 | 4561743 | 0.975 | 36384 | 113.567 |
| 1987-101 | 290857 | 0.968 | 12644 | 13.926 |
| 2069-101 | 3869190 | 0.978 | 36932 | 93.812 |
| 2093-101 | 2700416 | 0.98 | 37549 | 66.224 |
| 2102-101 | 2219077 | 0.974 | 34680 | 55.023 |
| 2172-15 | 980024 | 0.938 | 19945 | 34.482 |
| 2172-2 | 2497911 | 0.943 | 28183 | 75.962 |
| 2181-101 | 4189761 | 0.974 | 39622 | 100.017 |

| | | | | |
|---|---|---|---|---|
| 2186-101 | 2449823 | 0.979 | 32297 | 64.939 |
| 2401-101 | 2238459 | 0.972 | 34308 | 55.845 |
| 2412-101 | 555326 | 0.961 | 16565 | 21.261 |
| 251318-101 | 2181450 | 0.977 | 29419 | 59.609 |
| 251319-101 | 977057 | 0.977 | 23040 | 30.38 |
| 251320-101 | 3661717 | 0.974 | 40335 | 85.945 |
| 251321-101 | 2735553 | 0.978 | 27966 | 77.639 |
| 2544-101 | 4029122 | 0.977 | 36496 | 100.314 |
| 2576-101 | 2840786 | 0.971 | 32821 | 73.02 |
| 2578-101 | 1676259 | 0.974 | 28117 | 46.642 |
| 2649-101 | 420490 | 0.964 | 14742 | 17.394 |
| 2650-101 | 229441 | 0.981 | 9556 | 13.814 |
| 2725-101 | 4544274 | 0.978 | 37961 | 112.358 |
| 2832-101 | 3536608 | 0.973 | 38029 | 86.212 |
| 2839-101 | 4515964 | 0.969 | 36132 | 116.373 |
| 2852-101 | 1051515 | 0.973 | 22886 | 32.64 |
| 2854-101 | 1761127 | 0.979 | 26764 | 50.474 |
| 2857-101 | 2620451 | 0.98 | 37788 | 63.593 |
| 2915-101 | 3738875 | 0.976 | 34888 | 95.234 |
| 2933-101 | 73476 | 0.954 | 6306 | 6.1 |
| 2983-101 | 4009330 | 0.978 | 34339 | 104.643 |
| 3123-101 | 1426303 | 0.971 | 29396 | 37.907 |
| 365909-101 | 1356728 | 0.98 | 27700 | 37.888 |
| 365910-101 | 3475812 | 0.982 | 35251 | 87.426 |
| 365911-101 | 17586 | 0.94 | 3173 | 2.728 |
| 365912-101 | 2690411 | 0.98 | 28908 | 75.043 |
| 365914-101 | 2912501 | 0.978 | 39856 | 68.799 |
| 365915-101 | 2507336 | 0.98 | 36527 | 61.872 |
| 365916-101 | 1756143 | 0.978 | 31283 | 46.149 |
| 365917-101 | 1106543 | 0.98 | 22047 | 35.6 |
| 365918-101 | 3367378 | 0.979 | 42736 | 76.855 |
| 365958-101 | 4835275 | 0.975 | 37191 | 118.94 |
| 365960-101 | 2534529 | 0.977 | 30820 | 67.633 |
| 365961-101 | 2291525 | 0.978 | 29046 | 64.786 |
| 365962-101 | 3773949 | 0.975 | 36760 | 92.937 |
| 365964-101 | 3201558 | 0.976 | 36215 | 79.441 |
| 365965-101 | 3722221 | 0.978 | 28182 | 104.862 |
| 365966-101 | 1822739 | 0.979 | 25301 | 55.208 |
| 3778-5 | 2004880 | 0.925 | 21279 | 74.21 |
| 3778-9 | 1646601 | 0.927 | 20385 | 63.181 |

| | | | | |
|---|---|---|---|---|
| 379021-101 | 1778550 | 0.977 | 28344 | 49.177 |
| 379023-101 | 2888964 | 0.977 | 34130 | 73.825 |
| 379024-101 | 1718608 | 0.973 | 25672 | 50.495 |
| 379025-101 | 4022951 | 0.978 | 34063 | 103.335 |
| 379026-101 | 4340700 | 0.98 | 38470 | 104.786 |
| 379059-101 | 2421589 | 0.981 | 31746 | 64.709 |
| 390519-101 | 1666679 | 0.977 | 29865 | 44.378 |
| 390688-101 | 1752668 | 0.978 | 30987 | 46.157 |
| 390689-101 | 4512953 | 0.977 | 38270 | 110.085 |
| 390690-101 | 2646066 | 0.976 | 10851 | 150.443 |
| 390691-101 | 3381952 | 0.976 | 36319 | 83.565 |
| 390695-101 | 423532 | 0.976 | 14109 | 18.452 |
| 390696-101 | 3692047 | 0.976 | 36228 | 90.892 |
| 390699-101 | 2130632 | 0.975 | 31947 | 55.986 |
| 390700-101 | 3748237 | 0.975 | 34294 | 95.713 |
| 390701-101 | 2286853 | 0.976 | 35498 | 56.239 |
| 390705-101 | 2541073 | 0.976 | 38360 | 60.362 |
| 390706-101 | 5667961 | 0.979 | 42438 | 132.915 |
| 390707-101 | 2150065 | 0.977 | 32006 | 55.456 |
| 390716-101 | 649115 | 0.978 | 23213 | 20.37 |
| 390718-101 | 2263338 | 0.976 | 34473 | 55.94 |
| 390721-101 | 1990569 | 0.978 | 32194 | 52.027 |
| 390722-101 | 2426132 | 0.978 | 30980 | 64.961 |
| 390728-101 | 4222460 | 0.978 | 36856 | 105.316 |
| 390748-102 | 1619440 | 0.978 | 28684 | 44.355 |
| 407533-101 | 1313574 | 0.973 | 29425 | 36.275 |
| 407534-101 | 3650109 | 0.976 | 36673 | 88.856 |
| 407535-103 | 2199289 | 0.976 | 33381 | 56.649 |
| 407537-101 | 3609456 | 0.975 | 32089 | 94.558 |
| 407538-101 | 2500804 | 0.972 | 33091 | 64.03 |
| 407544-101 | 1986374 | 0.972 | 31221 | 51.989 |
| 407550-101 | 4712208 | 0.978 | 36252 | 118.624 |
| 407551-101 | 2952304 | 0.969 | 33357 | 75.274 |
| 407552-101 | 2026851 | 0.978 | 25959 | 59.894 |
| 407553-101 | 3324274 | 0.977 | 33136 | 86.48 |
| 407554-101 | 1600623 | 0.974 | 30346 | 42.51 |
| 4117A-103 | 2270802 | 0.925 | 21943 | 82.778 |
| 4117A-31 | 2189807 | 0.927 | 24388 | 73.222 |
| 503517-101 | 41587 | 0.977 | 6038 | 3.769 |
| 503519-101 | 2150172 | 0.976 | 30221 | 57.298 |

| | | | | |
|---|---|---|---|---|
| 503521-101 | 6484748 | 0.979 | 45894 | 147.141 |
| 503522-101 | 2816838 | 0.975 | 35615 | 70.381 |
| 503523-101 | 2419634 | 0.973 | 31376 | 63.588 |
| 503524-101 | 1960823 | 0.982 | 30638 | 53.171 |
| MG101-1 | 725195 | 0.928 | 18335 | 25.537 |
| MG103-1 | 977295 | 0.96 | 18436 | 35.374 |
| MG103-2 | 872069 | 0.941 | 19843 | 29.093 |
| MG104-1 | 1057938 | 0.958 | 11116 | 58.649 |
| MG105-1R | 802728 | 0.958 | 20614 | 25.944 |
| MG105-2 | 1251346 | 0.955 | 22039 | 38.886 |
| MG105-3 | 913195 | 0.957 | 14281 | 37.933 |
| MG105-4 | 419832 | 0.941 | 14741 | 16.625 |
| MG105-5 | 1097887 | 0.95 | 18938 | 37.605 |
| MG105-6 | 620727 | 0.928 | 15240 | 24.131 |
| MG105-7 | 1065864 | 0.94 | 22244 | 32.856 |
| MG105-8 | 1669040 | 0.946 | 21116 | 53.846 |
| MG106-1 | 1048647 | 0.935 | 21094 | 33.885 |
| MG106-2 | 2067945 | 0.951 | 24944 | 61.98 |
| MG106-3 | 1508324 | 0.919 | 25277 | 42.628 |
| MG107-1 | 1020866 | 0.949 | 20784 | 33.154 |
| MG107-10 | 1371695 | 0.95 | 23053 | 41.631 |
| MG107-2 | 1013620 | 0.954 | 19659 | 34.407 |
| MG107-3 | 1570291 | 0.943 | 23608 | 46.753 |
| MG107-4 | 1191986 | 0.947 | 22672 | 36.376 |
| MG107-5 | 1233479 | 0.949 | 23085 | 37.537 |
| MG107-6 | 1400747 | 0.951 | 24138 | 41.631 |
| MG107-7 | 1509667 | 0.943 | 25277 | 42.924 |
| MG107-8 | 817534 | 0.944 | 22834 | 24.648 |
| MG107-9 | 1247038 | 0.948 | 23293 | 37.524 |
| MG109-1 | 487019 | 0.934 | 16752 | 17.768 |
| MG109-2 | 1366389 | 0.922 | 23161 | 40.268 |
| MG110-1 | 1495047 | 0.944 | 19625 | 50.782 |
| MG111-1 | 1184637 | 0.951 | 22352 | 36.789 |
| MG111-10 | 1062784 | 0.923 | 19524 | 35.003 |
| MG111-11 | 971504 | 0.938 | 18317 | 34.064 |
| MG111-12 | 941272 | 0.932 | 16856 | 34.805 |
| MG111-13 | 1611403 | 0.9 | 10274 | 95.883 |
| MG111-14 | 652155 | 0.932 | 13871 | 27.868 |
| MG111-15 | 1149410 | 0.921 | 20600 | 37.079 |
| MG111-2 | 1319828 | 0.956 | 21336 | 42.519 |

| | | | | |
|---|---|---|---|---|
| MG111-3 | 650475 | 0.938 | 19195 | 21.732 |
| MG111-4 | 1467335 | 0.951 | 24297 | 43.762 |
| MG111-5 | 975986 | 0.939 | 18848 | 33.374 |
| MG111-6 | 840759 | 0.934 | 17313 | 30.492 |
| MG111-7 | 1203553 | 0.948 | 18638 | 42.454 |
| MG111-8 | 1160326 | 0.941 | 17337 | 42.655 |
| MG111-9 | 1297324 | 0.929 | 20208 | 42.701 |
| MG113-1 | 571968 | 0.772 | 8039 | 33.615 |
| MG113-10 | 1244608 | 0.938 | 24388 | 35.874 |
| MG113-2 | 653093 | 0.944 | 15521 | 25.727 |
| MG113-3R | 300687 | 0.943 | 13535 | 12.897 |
| MG113-4R | 843535 | 0.937 | 18422 | 28.902 |
| MG113-5 | 575110 | 0.935 | 16909 | 20.961 |
| MG113-6R | 2593997 | 0.944 | 25708 | 74.595 |
| MG113-7 | 850539 | 0.943 | 18776 | 28.988 |
| MG113-8 | 2269982 | 0.952 | 23278 | 68.572 |
| MG113-9 | 1278430 | 0.935 | 22422 | 38.587 |
| MG114-1 | 1611257 | 0.943 | 25513 | 45.471 |
| MG114-10 | 1796885 | 0.95 | 25910 | 51.116 |
| MG114-11R | 1613402 | 0.944 | 24758 | 46.904 |
| MG114-12 | 1507046 | 0.935 | 21076 | 47.935 |
| MG114-13 | 1342628 | 0.94 | 19642 | 44.421 |
| MG114-14 | 1317485 | 0.947 | 19916 | 43.755 |
| MG114-15 | 1591814 | 0.944 | 21411 | 50.567 |
| MG114-2 | 1300896 | 0.944 | 12767 | 60.66 |
| MG114-3 | 1723271 | 0.944 | 11520 | 89.564 |
| MG114-4 | 1603043 | 0.943 | 22800 | 48.25 |
| MG114-5 | 1228445 | 0.936 | 20553 | 39.003 |
| MG114-6 | 762486 | 0.937 | 17455 | 27.083 |
| MG114-7 | 1141877 | 0.942 | 16898 | 41.686 |
| MG114-8 | 1205965 | 0.947 | 24169 | 35.389 |
| MG114-9 | 1141136 | 0.955 | 15419 | 45.178 |
| MG115-1 | 1477117 | 0.938 | 23093 | 44.483 |
| MG115-10 | 973572 | 0.944 | 20389 | 32.138 |
| MG115-11 | 1231265 | 0.941 | 24113 | 36.387 |
| MG115-12 | 1567784 | 0.942 | 24255 | 46.785 |
| MG115-13 | 2586243 | 0.963 | 21691 | 85.989 |
| MG115-14 | 1941036 | 0.957 | 27676 | 53.886 |
| MG115-15 | 2145739 | 0.961 | 30369 | 57.138 |
| MG115-16 | 2070350 | 0.961 | 28224 | 57.151 |

| | | | | |
|---|---|---|---|---|
| MG115-17R | 1726535 | 0.952 | 24777 | 50.676 |
| MG115-18 | 1979915 | 0.963 | 24131 | 60.335 |
| MG115-19 | 2262119 | 0.97 | 17555 | 85.434 |
| MG115-2 | 1106841 | 0.926 | 24626 | 31.365 |
| MG115-20 | 2150328 | 0.961 | 28516 | 58.657 |
| MG115-3 | 1533423 | 0.938 | 24524 | 44.596 |
| MG115-4 | 1800051 | 0.948 | 23123 | 54.73 |
| MG115-5 | 1345188 | 0.932 | 23577 | 39.921 |
| MG115-6 | 922730 | 0.931 | 18308 | 32.439 |
| MG115-7 | 1383340 | 0.938 | 23713 | 40.832 |
| MG115-8 | 685980 | 0.942 | 17987 | 24.383 |
| MG115-9 | 856138 | 0.943 | 22300 | 26.311 |
| MG116-1 | 2620413 | 0.955 | 27302 | 73.4 |
| MG116-10 | 1474196 | 0.959 | 23858 | 44.53 |
| MG116-11 | 1157341 | 0.959 | 24159 | 34.504 |
| MG116-12 | 1533917 | 0.964 | 24423 | 46.12 |
| MG116-2 | 1364474 | 0.945 | 23098 | 40.972 |
| MG116-3 | 1327137 | 0.954 | 20642 | 44.534 |
| MG116-4 | 1621290 | 0.941 | 23231 | 48.327 |
| MG116-5 | 1527120 | 0.967 | 24202 | 45.724 |
| MG116-6 | 1905794 | 0.96 | 27432 | 52.444 |
| MG116-7 | 2153685 | 0.968 | 23252 | 67.593 |
| MG116-8 | 1782314 | 0.965 | 24634 | 52.398 |
| MG116-9 | 790971 | 0.957 | 18616 | 27.702 |
| MG117-1 | 2809514 | 0.961 | 28967 | 77.132 |
| MG117-10 | 1502902 | 0.958 | 26639 | 42.204 |
| MG117-11 | 1923532 | 0.95 | 28226 | 52.276 |
| MG117-12 | 1211097 | 0.959 | 22834 | 37.889 |
| MG117-2 | 3476301 | 0.96 | 30638 | 92.095 |
| MG117-3 | 1632177 | 0.964 | 24771 | 48.427 |
| MG117-4 | 2307685 | 0.956 | 26581 | 65.874 |
| MG117-5 | 1610157 | 0.954 | 24949 | 46.59 |
| MG117-6 | 1206813 | 0.951 | 21453 | 38.663 |
| MG117-7 | 1801874 | 0.946 | 26204 | 51.04 |
| MG117-8 | 1715525 | 0.953 | 24281 | 51.474 |
| MG117-9 | 2179544 | 0.958 | 26274 | 62.452 |
| MG118-1 | 1103683 | 0.941 | 19515 | 37.384 |
| MG118-2 | 1269249 | 0.955 | 22429 | 40.491 |
| MG118-3 | 1445480 | 0.956 | 21846 | 46.781 |
| MG118-4 | 1301649 | 0.954 | 23723 | 39.913 |

| | | | | |
|---|---|---|---|---|
| MG119-1 | 1354403 | 0.945 | 23381 | 41.215 |
| MG120-1 | 1317924 | 0.953 | 22436 | 41.413 |
| MG120-10 | 1063381 | 0.952 | 17837 | 38.961 |
| MG120-11-1 | 1041049 | 0.943 | 18670 | 36.437 |
| MG120-11-2 | 1009798 | 0.97 | 20749 | 33.949 |
| MG120-13 | 982881 | 0.959 | 20783 | 31.933 |
| MG120-15 | 1270578 | 0.964 | 23248 | 39.201 |
| MG120-17 | 2247542 | 0.967 | 23367 | 70.732 |
| MG120-18 | 2030229 | 0.967 | 24802 | 61.665 |
| MG120-19 | 2262015 | 0.973 | 13697 | 106.883 |
| MG120-2 | 886664 | 0.944 | 19009 | 30.448 |
| MG120-20 | 2718318 | 0.968 | 18435 | 104.637 |
| MG120-21 | 2427974 | 0.96 | 19247 | 85.343 |
| MG120-22 | 1318297 | 0.923 | 16698 | 49.39 |
| MG120-23 | 1374128 | 0.97 | 12017 | 72.572 |
| MG120-24 | 2036867 | 0.966 | 19389 | 72.398 |
| MG120-25R | 1643420 | 0.968 | 21039 | 54.345 |
| MG120-26 | 1708398 | 0.97 | 19285 | 60.138 |
| MG120-27 | 1396758 | 0.959 | 19094 | 49.861 |
| MG120-28 | 2766644 | 0.967 | 21790 | 91.26 |
| MG120-29 | 2761259 | 0.965 | 24255 | 85.876 |
| MG120-3 | 825272 | 0.903 | 18024 | 28.046 |
| MG120-30 | 2487753 | 0.956 | 21644 | 80.888 |
| MG120-31 | 2521148 | 0.952 | 15268 | 111.736 |
| MG120-32 | 3240975 | 0.966 | 25778 | 95.885 |
| MG120-33 | 2757052 | 0.964 | 25710 | 81.585 |
| MG120-34 | 1868699 | 0.963 | 24014 | 57.217 |
| MG120-35 | 1630391 | 0.961 | 12558 | 85.032 |
| MG120-4 | 1228342 | 0.949 | 18728 | 43.217 |
| MG120-5 | 935242 | 0.946 | 13198 | 42.036 |
| MG120-6 | 1011413 | 0.933 | 18383 | 35.368 |
| MG120-7 | 1012083 | 0.941 | 18590 | 35.337 |
| MG120-8 | 1593234 | 0.948 | 20843 | 53.045 |
| MG120-9 | 1572342 | 0.948 | 19438 | 54.613 |
| MG121-1 | 1945828 | 0.973 | 19266 | 69.698 |
| MG122-1 | 2686292 | 0.976 | 12638 | 144.7 |
| MG122-2 | 4586924 | 0.978 | 23482 | 150.272 |
| MG124-1R | 1017508 | 0.968 | 17320 | 37.994 |
| MG124-2 | 955443 | 0.954 | 20026 | 31.856 |
| MG124-3R | 2370949 | 0.969 | 24067 | 72.399 |

| | | | | |
|---|---|---|---|---|
| MG124-4R | 907607 | 0.971 | 19503 | 31.376 |
| MG125-1 | 2535112 | 0.964 | 24049 | 76.8 |
| MG125-10 | 2150726 | 0.964 | 27549 | 60.85 |
| MG125-2 | 3231953 | 0.973 | 25269 | 98.059 |
| MG125-3 | 3279297 | 0.974 | 23299 | 105.228 |
| MG125-4 | 4015968 | 0.958 | 25993 | 118.148 |
| MG125-5 | 1402070 | 0.957 | 23665 | 42.37 |
| MG125-6 | 1569825 | 0.958 | 25790 | 45.052 |
| MG125-7 | 2363601 | 0.963 | 27347 | 66.782 |
| MG125-8 | 2594420 | 0.962 | 28513 | 72.106 |
| MG125-9 | 1980450 | 0.961 | 26345 | 57.123 |
| MG126-1 | 1505836 | 0.963 | 25147 | 45.062 |
| MG126-2 | 1631100 | 0.973 | 14470 | 71.207 |
| MG126-3 | 1551656 | 0.966 | 19077 | 54.742 |
| MG126-4 | 1382566 | 0.958 | 23114 | 43.06 |
| MG126-5 | 1190730 | 0.93 | 24328 | 34.349 |
| MG127-1 | 707240 | 0.963 | 19487 | 24.347 |
| MG127-2 | 1420116 | 0.965 | 23195 | 44.543 |
| MG128-1 | 1454426 | 0.965 | 24074 | 44.481 |

**Table S5**: Summary of sequence and genotype filters

| Filter # | Filter applied | Script calls | # of SNPs | Used for |
|---|---|---|---|---|
| Initial Stacks calls | Internal Stacks quality filters | Stacks ref_map | | - |
| 1 | Remove stacks present in less than 80% of samples | Stacks populations -R 0.8 | 49,580 | Diversity and divergence*†; introgression HMM*†; *Treemix*† |
| 2 | Remove sites supported by less than 8 reads | vcftools –minDP –max-missing-count | 11,297 | - |
| 3 | Remove individuals with > 60% missing data | vcftools –remove-indv | 11,297 | D-stats |
| 4 | Remove sites in high LD ($r^2 > 0.7$) | bcftools +prune | 5,767 | *fastStructure; NewHybrids; RAxML; δaδi* |

* Performed on Stacks assembled haplotypes, not SNPs.
† Individuals with missing data also removed

**Table S6**: Full likelihoods for *NewHybrids* classifications of CHSxPIM admixture on Santa Cruz. MCMC was run for 6,000 steps. The most likely classification for each individual is shown in bold.

| Individual | Pop | CHS | PIM | F1 | F2 | BC_CHS | BC_PIM |
|---|---|---|---|---|---|---|---|
| MG113-10 | PIM | **1.00** | 0.00 | 0.00 | 0.00 | 0.00 | 0.00 |
| MG113-2 | PIM | **1.00** | 0.00 | 0.00 | 0.00 | 0.00 | 0.00 |
| MG113-3R | PIM | **1.00** | 0.00 | 0.00 | 0.00 | 0.00 | 0.00 |
| MG113-4R | PIM | **1.00** | 0.00 | 0.00 | 0.00 | 0.00 | 0.00 |
| MG113-5 | PIM | **1.00** | 0.00 | 0.00 | 0.00 | 0.00 | 0.00 |
| MG113-6R | PIM | **1.00** | 0.00 | 0.00 | 0.00 | 0.00 | 0.00 |
| MG113-7 | PIM | **1.00** | 0.00 | 0.00 | 0.00 | 0.00 | 0.00 |
| MG113-8 | PIM | **1.00** | 0.00 | 0.00 | 0.00 | 0.00 | 0.00 |
| MG113-9 | PIM | **1.00** | 0.00 | 0.00 | 0.00 | 0.00 | 0.00 |
| MG115-10 | CHS | 0.00 | **1.00** | 0.00 | 0.00 | 0.00 | 0.00 |
| MG115-11 | CHS | 0.00 | **1.00** | 0.00 | 0.00 | 0.00 | 0.00 |
| MG115-12 | CHS | 0.00 | **1.00** | 0.00 | 0.00 | 0.00 | 0.00 |
| MG115-13 | CHS | 0.00 | **1.00** | 0.00 | 0.00 | 0.00 | 0.00 |
| MG115-14 | CHS | 0.00 | **1.00** | 0.00 | 0.00 | 0.00 | 0.00 |
| MG115-15 | CHS | 0.00 | **1.00** | 0.00 | 0.00 | 0.00 | 0.00 |
| MG115-16 | ADMX | 0.00 | **1.00** | 0.00 | 0.00 | 0.00 | 0.00 |
| MG115-17R | ADMX | 0.00 | **1.00** | 0.00 | 0.00 | 0.00 | 0.00 |
| MG115-18 | CHS | 0.00 | **1.00** | 0.00 | 0.00 | 0.00 | 0.00 |
| MG115-19 | CHS | **1.00** | 0.00 | 0.00 | 0.00 | 0.00 | 0.00 |
| MG115-1 | CHS | 0.00 | **1.00** | 0.00 | 0.00 | 0.00 | 0.00 |
| MG115-20 | ADMX | 0.00 | 0.00 | 0.00 | **1.00** | 0.00 | 0.00 |
| MG115-2 | CHS | 0.00 | **1.00** | 0.00 | 0.00 | 0.00 | 0.00 |
| MG115-3 | ADMX | 0.00 | 0.00 | 0.00 | **1.00** | 0.00 | 0.00 |
| MG115-4 | CHS | 0.00 | **1.00** | 0.00 | 0.00 | 0.00 | 0.00 |
| MG115-5 | ADMX | 0.00 | 0.00 | **1.00** | 0.00 | 0.00 | 0.00 |
| MG115-6 | CHS | 0.00 | **1.00** | 0.00 | 0.00 | 0.00 | 0.00 |
| MG115-7 | ADMX | 0.00 | 0.00 | **1.00** | 0.00 | 0.00 | 0.00 |
| MG115-8 | CHS | 0.00 | **1.00** | 0.00 | 0.00 | 0.00 | 0.00 |
| MG115-9 | CHS | 0.00 | **1.00** | 0.00 | 0.00 | 0.00 | 0.00 |

**Table S7**: Full likelihoods for *NewHybrids* classifications of CHSxGAL admixture on Isabela. MCMC was run for 6,000 steps. The most likely classification for each individual is shown in bold.

| Individual | Pop | CHS | PIM | F1 | F2 | BC_CHS | BC_PIM |
|---|---|---|---|---|---|---|---|
| MG120-13 | ADMX | 0.00000 | 0.00000 | 0.00000 | **1.00000** | 0.00000 | 0.00000 |
| MG120-15 | ADMX | **0.99983** | 0.00000 | 0.00000 | 0.00017 | 0.00000 | 0.00000 |
| MG120-19 | ADMX | 0.00000 | **0.99983** | 0.00000 | 0.00017 | 0.00000 | 0.00000 |
| MG120-23 | ADMX | 0.00000 | 0.00000 | 0.00000 | 0.00017 | 0.00000 | **0.99982** |
| MG120-28 | ADMX | 0.00000 | 0.00000 | 0.07191 | 0.04213 | 0.00000 | **0.88596** |
| MG120-33 | ADMX | 0.00000 | 0.00000 | 0.00000 | **0.99986** | 0.00000 | 0.00014 |
| MG120-34 | ADMX | 0.00000 | 0.00000 | 0.00003 | **0.99995** | 0.00002 | 0.00000 |
| MG120-11-2 | CHS | **0.99983** | 0.00000 | 0.00000 | 0.00017 | 0.00000 | 0.00000 |
| MG120-17 | CHS | **0.99983** | 0.00000 | 0.00000 | 0.00017 | 0.00000 | 0.00000 |
| MG120-18 | CHS | **0.99983** | 0.00000 | 0.00000 | 0.00017 | 0.00000 | 0.00000 |
| MG120-20 | CHS | **0.99983** | 0.00000 | 0.00000 | 0.00017 | 0.00000 | 0.00000 |
| MG120-22 | CHS | **0.99983** | 0.00000 | 0.00000 | 0.00017 | 0.00000 | 0.00000 |
| MG120-24 | CHS | 0.01289 | 0.00000 | 0.00000 | **0.75656** | 0.00000 | 0.23055 |
| MG120-27 | CHS | 0.00000 | 0.00000 | 0.19540 | 0.00655 | 0.00000 | **0.79804** |
| MG120-29 | CHS | **0.99983** | 0.00000 | 0.00000 | 0.00017 | 0.00000 | 0.00000 |
| MG120-31 | CHS | **0.99983** | 0.00000 | 0.00000 | 0.00017 | 0.00000 | 0.00000 |
| MG120-10 | PIM | 0.00000 | **0.99983** | 0.00000 | 0.00017 | 0.00000 | 0.00000 |
| MG120-11-1 | PIM | 0.00000 | **0.99983** | 0.00000 | 0.00017 | 0.00000 | 0.00000 |
| MG120-21 | PIM | 0.00000 | **0.99983** | 0.00000 | 0.00017 | 0.00000 | 0.00000 |
| MG120-25R | PIM | 0.00000 | **0.99983** | 0.00000 | 0.00017 | 0.00000 | 0.00000 |
| MG120-26 | PIM | 0.00000 | 0.00000 | 0.00000 | 0.00843 | **0.99157** | 0.00000 |
| MG120-2 | PIM | 0.00000 | **0.99983** | 0.00000 | 0.00017 | 0.00000 | 0.00000 |
| MG120-1 | PIM | 0.00000 | **0.99983** | 0.00000 | 0.00017 | 0.00000 | 0.00000 |
| MG120-30 | PIM | 0.00000 | **0.99983** | 0.00000 | 0.00017 | 0.00000 | 0.00000 |
| MG120-32 | PIM | 0.00000 | **0.99983** | 0.00000 | 0.00017 | 0.00000 | 0.00000 |
| MG120-3 | PIM | 0.00000 | **0.99983** | 0.00000 | 0.00017 | 0.00000 | 0.00000 |
| MG120-4 | PIM | 0.00000 | **0.99983** | 0.00000 | 0.00017 | 0.00000 | 0.00000 |
| MG120-5 | PIM | 0.00000 | **0.99983** | 0.00000 | 0.00017 | 0.00000 | 0.00000 |
| MG120-6 | PIM | 0.00000 | **0.99983** | 0.00000 | 0.00017 | 0.00000 | 0.00000 |
| MG120-7 | PIM | 0.00000 | **0.99983** | 0.00000 | 0.00017 | 0.00000 | 0.00000 |
| MG120-8 | PIM | 0.00000 | **0.99983** | 0.00000 | 0.00017 | 0.00000 | 0.00000 |
| MG120-9 | PIM | 0.00000 | **0.99983** | 0.00000 | 0.00017 | 0.00000 | 0.00000 |

**Table S8**: *NewHybrids* MCMC summary. Only the most likely genotype category classifications are shown. Refer to Table S5 for alternative class probabilities.

| Population A | Population B | Type | Classification | | | |
|---|---|---|---|---|---|---|
| | | | Pure | $F_1$ | $F_2$ | BC |
| MG115 | MG113 | CHS-PIM | 25 | 2 | 2 | 0 |
| MG120C | MG120G | CHS-GAL | 24 | 0 | 4 | 4 |

**Table S9**: Likelihoods for different *Treemix* runs

| m | ln(L) |
|---|---|
| 0 | 157.79953 |
| 1 | 350.8362 |
| 2 | 374.0593 |
| 3 | 381.47799 |
| 4 | 389.98895 |
| 5 | 393.66234 |
| 6 | 395.08137 |
| 7 | 396.37992 |
| 8 | 396.8767 |

**Table S10**: Mean Fst between focal island populations and mainland groups.

| | **Peru** | **Ecu** |
|---|---|---|
| **MG105** | 0.351069 | 0.193202 |
| **MG106** | 0.335434 | 0.25599 |
| **MG107** | 0.338202 | 0.172407 |
| **MG111** | 0.357811 | 0.21111 |
| **MG113** | 0.329355 | 0.169055 |
| **MG114** | 0.348367 | 0.188794 |
| **MG115** | 0.362342 | 0.298977 |
| **MG117** | 0.342618 | 0.182225 |
| **MG120_G** | 0.387008 | 0.347495 |
| **MG120_C** | 0.412477 | 0.370005 |
| **MG120_GC** | 0.380804 | 0.329513 |
| **MG125** | 0.308738 | 0.130834 |
| **MG126** | 0.33319 | 0.190727 |

**Table S11**: Summary of inferred introgression blocks for population MG114 (polymorphic). Shaded groups indicate blocks that are likely the same age/the result of the same hybridization event based on break points.

| Block | Size (Mb) | Chr | Start (Mb) | Stop (Mb) | Count | Individuals | Type |
|---|---|---|---|---|---|---|---|
| 114_1A | 1.3 | 1 | 80 | 81.3 | 1 | MG114-11 | CHS/CHS |
| 114_1B | 1.2 | 1 | 80.1 | 81.3 | 1 | MG114-14 | CHS/CHS |
| 114_1C | 0.30 | 1 | 85.20 | 85.50 | 1 | MG114-13 | CHS/CHS |
| 114_1D | 6.8 | 1 | 16.0 | 22.8 | 1 | MG114-9 | CHS/CHS |
| 114_1E | 0.7 | 1 | 76.6 | 77.3 | 1 | MG114-9 | CHS/CHS |
| 114_2A | 3.60 | 2 | 36.9 | 40.1 | 1 | MG114-7 | CHS/CHS |
| 114_3A | 5.0 | 3 | 23.0 | 28.0 | 1 | MG114-12 | CHS/CHS |
| 114_3B | 3.1 | 3 | 4.8 | 7.9 | 1 | MG114-4 | CHS/CHS |
| 114_3C | 1.70 | 3 | 3.54 | 5.10 | 1 | MG114-11 | CHS/CHS |
| 114_3D* | 50.9 | 3 | 4.50 | 55.4 | 1 | MG114-1 | CHS/CHS |
| 114_3E* | 51.8 | 3 | 3.6 | 55.4 | 5 | MG114-13, MG114-2, MG114-6, MG114-3, MG114-14 | CHS/CHS |
| 114_4A | 8.70 | 4 | 8.40 | 17.10 | 1 | MG114-13 | CHS/CHS |
| 114_4B | 9.2 | 4 | 7.20 | 16.4 | 1 | MG114-3 | CHS/CHS |
| 114_4C | 8.0 | 4 | 8.4 | 16.4 | 1 | MG114-14 | CHS/CHS |
| 114_4D | 9.3 | 4 | 16.9 | 26.2 | 1 | MG114-11 | CHS/CHS |
| 114_4E | 4.8 | 4 | 43.0 | 48.8 | 1 | MG114-11 | CHS/CHS |
| 114_5A | 2.2 | 5 | 60.3 | 62.5 | 1 | MG114-3 | CHS/CHS |
| 114_6A† | 34.70 | 6 | 0.00 | 35.70 | 1 | MG114-9 | CHS/CHS |
| 114_6B† | 36.70 | 6 | 0.00 | 36.70 | 1 | MG114-4 | CHS/CHS |
| 114_6C† | 34.50 | 6 | 1.50 | 36.00 | 1 | MG114-1 | CHS/CHS |
| 114_6D | 4.30 | 6 | 1.1 | 25.7 | 1 | MG114-10 | CHS/CHS |
| 114_6E | 7.9 | 6 | 12.6 | 20.5 | 1 | MG114-12 | CHS/CHS |
| 114_6F | 5.4 | 6 | 29.7 | 35.1 | 1 | MG114-12 | CHS/CHS |
| 114_6G | 0.90 | 6 | 36.50 | 37.40 | 1 | MG114-7 | CHS/CHS |
| 114_6H | 19.7 | 6 | 7.0 | 26.7 | 1 | MG114-15 | CHS/CHS |
| 114_7A | 7.9 | 7 | 12.6 | 20.5 | 1 | MG114-3 | CHS/CHS |
| 114_7B | 5.4 | 7 | 29.7 | 35.1 | 1 | MG114-3 | CHS/CHS |
| 114_7C | 0.80 | 7 | 2.30 | 3.10 | 1 | MG114-3 | CHS/CHS |
| 114_9A | 0.60 | 9 | 72.3 | 72.9 | 1 | MG114-10 | CHS/CHS |
| 114_11A | 1.9 | 11 | 50.9 | 52.8 | 1 | MG114-3 | CHS/CHS |

*CHS chromosome 3 haplotype; †CHS chromosome 6 haplotype;

**Table S12**: Summary of inferred introgression blocks for population MG117 (polymorphic). Shaded groups indicate blocks that are likely the same age/the result of the same hybridization event based on break points.

| Block | Size (Mb) | Chr | Start (Mb) | Stop (Mb) | Count | Individuals | Type |
|---|---|---|---|---|---|---|---|
| 117_1A | 6.6 | 1 | 51.4 | 58.0 | 1 | MG117-9 | CHS/CHS |
| 117_1B | 0.7 | 1 | 76.6 | 77.3 | 1 | MG117-9 | CHS/CHS |
| 117_1C | 5.8 | 1 | 4.1 | 9.9 | 1 | MG117-12 | CHS/CHS |
| 117_1D | 4.2 | 1 | 4.1 | 8.3 | 1 | MG117-5 | CHS/CHS |
| 117_1E | 17 | 1 | 44 | 61 | 3 | MG117-12, MG117-2, MG117-5 | CHS/CHS |
| 117_1F | 10.1 | 1 | 5.7 | 15.8 | 1 | MG117-2 | CHS/CHS |
| 117_1G | 5.7 | 1 | 69.3 | 75.0 | 1 | MG117-2 | CHS/CHS |
| 117_1H | 33.8 | 1 | 31 | 64.8 | 1 | MG117-6 | CHS/CHS |
| 117_2A | 2.0 | 2 | 29.2 | 31.2 | 1 | MG117-2 | CHS/CHS |
| 117_2B | 3.9 | 2 | 28.5 | 32.4 | 1 | MG117-6 | CHS/CHS |
| 117_2C | 0.3 | 2 | 55.6 | 55.9 | 1 | MG117-2 | CHS/CHS |
| 117_3A | 1.6 | 3 | 66.2 | 67.8 | 2 | MG117-3, MG117-5 | CHS/CHS |
| 117_3B | 3.2 | 3 | 7.2 | 10.4 | 1 | MG117-12 | CHS/CHS |
| 117_3B | 2.3 | 3 | 8.1 | 10.4 | 1 | MG117-5 | CHS/CHS |
| 117_4A* | 2.2 | 4 | 64.4 | 66.6 | 1 | MG117-8 | CHS/CHS |
| 117_4B* | 2.9 | 4 | 63.7 | 66.6 | 2 | MG117-11, MG117-10 | CHS/CHS |
| 117_4C* | 1.9 | 4 | 64.7 | 66.6 | 1 | MG117-7 | CHS/CHS |
| 117_4D | 8 | 4 | 8.4 | 16.4 | 2 | MG117-12, MG117-5 | CHS/CHS |
| 117_6A† | 36.3 | 6 | 1.1 | 37.4 | 4 | MG117-8, MG117-11, MG117-7, MG117-10 | CHS/CHS |
| 117_6B† | 7.3 | 6 | 42.5 | 49.8 | 1 | MG117-8 | CHS/CHS |
| 117_6C† | 5.3 | 6 | 44.5 | 49.8 | 1 | MG117-11 | CHS/CHS |
| 117_6D† | 5.4 | 6 | 44.4 | 49.8 | 1 | MG117-7 | CHS/CHS |
| 117_6E† | 4.0 | 6 | 45.8 | 49.8 | 1 | MG117-10 | CHS/CHS |
| 117_6F | 6.7 | 6 | 43.1 | 49.8 | 2 | MG117-4, MG117-1 | **CHS/PIM** |
| 117_6G | 2.4 | 6 | 47.4 | 49.8 | 1 | MG117-6 | CHS/CHS |
| 117_6H | 5.1 | 6 | 32.2 | 37.3 | 1 | MG117-5 | CHS/CHS |
| 117_6I | 21.3 | 6 | 4.3 | 25.7 | 1 | MG117-6 | CHS/CHS |
| 117_7A | 1.2 | 7 | 5.2 | 6.4 | 1 | MG117-9 | CHS/CHS |
| 117_8A | 0.3 | 8 | 3.1 | 3.4 | 2 | MG117-5, MG117-12 | CHS/CHS |
| 117_9A | 0.6 | 9 | 72.3 | 72.9 | 2 | MG117-5, MG117-5 | CHS/CHS |
| 117_10A | 11.8 | 10 | 44.3 | 56.1 | 2 | MG117-3, MG117-2 | CHS/CHS |
| 117_10B | 1.3 | 10 | 59.2 | 60.5 | 1 | MG117-3 | CHS/CHS |
| 117_11A | 0.5 | 11 | 56.1 | 56.6 | 1 | MG117-3 | CHS/CHS |

| | | | | | | | |
|---|---|---|---|---|---|---|---|
| 117_11B | 30.9 | 11 | 14.8 | 45.7 | 1 | MG117-12 | CHS/CHS |
| 117_11C | 34.8 | 11 | 14.8 | 49.6 | 1 | MG117-5 | CHS/CHS |
| 117_11D | 0.6 | 11 | 50.9 | 51.5 | 1 | MG117-5 | CHS/CHS |

*CHS chromosome 4 haplotype; †CHS chromosome 6 haplotype

**Table S13:** Summary of inferred introgression blocks for population MG116 (fixed, red-fruited)

| Block | Size (Mb) | Chr | Start (Mb) | Stop (Mb) | Count | Individuals | Type |
|---|---|---|---|---|---|---|---|
| 116_1A | 4.6 | 1 | 4.8 | 9.4 | 1 | MG116-9 | CHS/CHS |
| 116_2A | 4.9 | 1 | 14.7 | 19.6 | 1 | MG116-9 | CHS/CHS |
| 116_2A | 20.4 | 2 | 0.4 | 20.8 | 1 | MG116-4 | CHS/CHS |
| 116_2B | 2.1 | 2 | 29.1 | 31.2 | 1 | MG116-4 | CHS/CHS |
| 116_2C | 1 | 2 | 39.9 | 40.9 | 1 | MG116-4 | CHS/CHS |
| 116_5A | 0.9 | 5 | 3.6 | 4.5 | 1 | MG116-6 | CHS/CHS |
| 116_6A | 1.1 | 6 | 45.6 | 46.7 | 1 | MG116-4 | CHS/CHS |
| 116_9A | 0.6 | 9 | 72.3 | 72.9 | 1 | MG116-6 | CHS/CHS |

**Table S14:** Population MG114 ancestry at carotenoid biosynthesis loci, as inferred by our HMM. The genomic location of each locus was determined based on the *Solanum lycopersicum* reference build SL3.0 and ITAG3.0 annotation. The association between ancestry and fruit color at PSY1 is significant based on a $\chi^2$ test of independence ($\chi^2 = 11.123$; df = 1; P = 0.00085). The association at LCY-B is not statistically significant ($\chi^2 = 2.934$; df = 1; P = 0.08673).

| Gene | | **ZDS** | **PSY1*** | **PDS** | **LCY-B** | **CYC-B** | **CCS** | **CRTISO** | **LCY-E** |
|---|---|---|---|---|---|---|---|---|---|
| Chromosome | | 1 | 3 | 3 | 4 | 6 | 8 | 10 | 12 |
| Position (bp) | | 88514122 | 4325334 | 70499494 | 11946753 | 45897927 | 63274891 | 62681526 | 2284379 |
| | **Fruit color** | **Ancestry** | | | | | | | |
| MG114-1 | Red | PIM | PIM | PIM | PIM | PIM | PIM | PIM | PIM |
| MG114-2 | Orange | PIM | CHS | PIM | PIM | PIM | PIM | PIM | PIM |
| MG114-3 | Orange | PIM | CHS | PIM | CHS | PIM | PIM | PIM | PIM |
| MG114-4 | Red | PIM | PIM | PIM | PIM | PIM | PIM | PIM | PIM |
| MG114-5 | Red | PIM | PIM | PIM | PIM | PIM | PIM | PIM | PIM |
| MG114-6 | Orange | PIM | CHS | PIM | PIM | PIM | PIM | PIM | PIM |
| MG114-7 | Red | PIM | PIM | PIM | PIM | PIM | PIM | PIM | PIM |
| MG114-8 | Red | PIM | PIM | PIM | PIM | PIM | PIM | PIM | PIM |
| MG114-9 | Red | PIM | PIM | PIM | PIM | PIM | PIM | PIM | PIM |
| MG114-10 | Red | PIM | PIM | PIM | PIM | PIM | PIM | PIM | PIM |
| MG114-11 | Orange | PIM | CHS | PIM | PIM | PIM | PIM | PIM | PIM |
| MG114-12 | Red | PIM | PIM | PIM | PIM | PIM | PIM | PIM | PIM |
| MG114-13 | Orange | PIM | CHS | PIM | CHS | PIM | PIM | PIM | PIM |
| MG114-14 | Orange | PIM | CHS | PIM | CHS | PIM | PIM | PIM | PIM |
| MG114-15 | Red | PIM | PIM | PIM | PIM | PIM | PIM | PIM | PIM |

**Table S15:** Population MG117 ancestry at carotenoid biosynthesis loci, as inferred by our HMM. The genomic location of each locus was determined based on the *Solanum lycopersicum* reference build SL3.0 and ITAG3.0 annotation. The association between ancestry and fruit color at CYC-B is significant based on a $\chi^2$ test of independence ($\chi^2 = 8.333$; df = 1; P = 0.00389). The association at LCY-B is not statistically significant ($\chi^2 = 0.6$; df = 1; P = 0.4386).

| Gene | | **ZDS** | **PSY1** | **PDS** | **LCY-B** | **CYC-B*** | **CCS** | **CRTISO** | **LCY-E** |
|---|---|---|---|---|---|---|---|---|---|
| Chromosome | | 1 | 3 | 3 | 4 | 6 | 8 | 10 | 12 |
| Position (bp) | | 88514122 | 4325334 | 70499494 | 11946753 | 45897927 | 63274891 | 62681526 | 2284379 |
| | **Fruit Color** | **Ancestry** | | | | | | | |
| MG117-1 | Orange | PIM | PIM | PIM | PIM | CHS | PIM | PIM | PIM |
| MG117-2 | Red | PIM | PIM | PIM | PIM | PIM | PIM | PIM | PIM |
| MG117-3 | Red | PIM | PIM | PIM | PIM | PIM | PIM | PIM | PIM |
| MG117-4 | Orange | PIM | PIM | PIM | PIM | CHS | PIM | PIM | PIM |
| MG117-5 | Red | PIM | PIM | PIM | CHS | PIM | PIM | PIM | PIM |
| MG117-6 | Red | PIM | PIM | PIM | PIM | PIM | PIM | PIM | PIM |
| MG117-7 | Orange | PIM | PIM | PIM | PIM | CHS | PIM | PIM | PIM |
| MG117-8 | Orange | PIM | PIM | PIM | PIM | CHS | PIM | PIM | PIM |
| MG117-9 | Red | PIM | PIM | PIM | PIM | PIM | PIM | PIM | PIM |
| MG117-10 | Orange | PIM | PIM | PIM | PIM | CHS | PIM | PIM | PIM |
| MG117-11 | Orange | PIM | PIM | PIM | PIM | CHS | PIM | PIM | PIM |
| MG117-12 | Red | PIM | PIM | PIM | CHS | PIM | PIM | PIM | PIM |

**Table S16**: Demographic model estimates for PIM population MG114 inferred using *δaδi* on the unmasked dataset (including introgressed regions). 95% CI values were obtained from 2,000 bootstrap replicates of the SFS. Each estimate is shown in rescaled units (rescaled by $N_{Ref}$ for $N_B$ and $N_F$; and by $2N_{Ref}$ for $T_B$ and $T_F$).

| Parameter | Optimum | Bootstrap Median | 95% CI |
|---|---|---|---|
| $N_B$ | 115.37 | 314.33 | 4.82 - 1444.46 |
| $N_F$ | 335.06 | 991.97 | 345.55 – 6644.64 |
| $T_B$ | 2125.97 | 928.21 | 109.74 – 8509.09 |
| $T_F$ | 52.59 | 917.73 | 35.77 – 16116.06 |
| F | 0.0008 | 0.0134 | $3.48 \times 10^{-9}$ – 0.53 |
